# MiRNA and phasiRNAs-mediated regulation of TIR-NBS-LRR defense genes in *Arabidopsis thaliana*

**DOI:** 10.1101/2020.03.02.972620

**Authors:** Diego López-Márquez, Ángel Del-Espino, Nieves López-Pagán, Edgar A. Rodríguez-Negrete, Ignacio Rubio-Somoza, Javier Ruiz-Albert, Eduardo R. Bejarano, Carmen R. Beuzón

## Abstract

Plants encode large numbers of intracellular immune receptors known as resistance (R) proteins or nucleotide-binding (NB) leucine-rich repeat (LRR) receptors (NLRs), involved in perception of pathogen-derived effectors and activation of immunity.

Here, we report a two-tiered regulatory network mediated by microRNA and secondary phased small RNAs (phasiRNA) that targets the silencing of dozens of NLR genes encoding yet uncharacterized members of the Toll/interleukin-1 (TIR)-NBS-LRR (TNLs) subfamily in *Arabidopsis*. We show that miR825-5p downregulates expression of *Arabidopsis AT5G38850* gene (renamed as *microRNA-silenced TNL 1 or MIST1*) by targeting the sequence coding for a highly conserved functional amino acid motif (TIR2) within the TIR domain of the receptor. Further, we show that *MIST1* functions as a regulatory hub, since miRNA825-5p triggers RDR6-mediated processing of *MIST1* transcripts, to generate *trans*-acting phasiRNAs that in turn target, a wide network of TNL genes for gene silencing. Regulation through *MIST1* affects disease resistance against the model bacterial pathogen *Pseudomonas syringae*, since altered levels of miRNA825-5p lead to changes in *Arabidopsis* ability to establish basal defenses against this pathogen. MiR825-5p is expressed in unchallenged adult leaves and its production is down regulated in response to PAMPs such as bacterial flagellin but also fungal chitin.

## Introduction

Plants possess complex immune systems that effectively protect them from the majority of pathogens present in the environment. The correct functioning of these systems rely on a battery of cell surface and intracellular receptors that alert plants of incoming threats, through the perception of pathogen-associated molecules or activities, and activate a cascade of defense responses capable of hindering disease development [1–3]. Immune receptors within the cell surface mediate perception of conserved molecules collectively known as pathogen-associated molecular patterns (PAMPs), and signal the activation of basal resistance, also known as PAMP-triggered immunity (PTI) [4]. Intracellular immune receptors detect either the presence inside the cell, or the perturbations caused within it by pathogen virulence factors (effectors), and activate a rapid defense response known as effector-triggered immunity (ETI) [1]. ETI reinstates and enhances PTI and is often associated to the activation of localized programmed cell death known as the hypersensitive response or HR [1]. Most of these intracellular receptors belong to a large family of proteins known as NOD-like receptor (NLR) proteins, which constitute the largest type of resistance (R) proteins, and are characterized by a multi-domain structure including a variable N-terminal domain, a nucleotide-binding domain (NB-ARC) and a leucine-rich repeat domain (LRR), hence their also being known as NBS-LRR proteins [1,3,5]. Most NLRs can be classified into two major families on the basis on their type of N-terminal domain: those displaying a Toll/interleukin-1 (TIR) domain, known as TNLs (TIR-NBS-LRR), and those with a domain that resembles a coiled-coil (CC) domain, generally known as CNLs (CC-NBS-LRR) [6]. TNLs and CNLs engage the ETI machinery through different key regulators of plant defense. ETI signaling *via* CNLs relies on NDR1 (non-race-specific disease resistance 1), while ETI signaling *via* TNLs requires the function of EDS1 (enhanced disease susceptibility 1), a lipase-like protein that conveys all TNL resistance outputs[7–10]. Additionally, TNL-mediated ETI responses could enhance basal immunity through the interaction of EDS1 with PAD4 (Phytoalexin deficient 4) [11–14].

Expression of NLR-resistance pathways is tightly controlled in the absence of the pathogen since constitutive activation can give rise to deleterious effects, *i*.*e*. mutants in negative regulators show developmental defects, while increased levels of NLRs can lead to activation of defense-related phenotypes such as cell wall modifications, production of reactive oxygen species (ROS) or spontaneous activation of the HR [15–20]. Thus, NLR protein production is down regulated by several mechanisms at transcriptional, post-transcriptional, translational, and post-translational levels. Small RNAs (sRNAs) are among the regulators that control NLR production acting at transcriptional or post-transcriptional level. Regarding post-transcriptional regulation, 21-22 nucleotide-long (21-22 nt) bind to their target mRNA by base pairing, and reduce the expression of the mRNA-encoded protein either by altering mRNA stability or its translation, through the action of proteins from the Argonaute family recruited to this purpose, a process known as post-transcriptional gene silencing (PTGS) [21, 22]. Two types of sRNAs, 21-22 nt microRNAs (miRNAs) and 21 nt small interfering RNAs (siRNAs), function as suppressors of NLR-encoding mRNAs [23]. PTGS of NLR expression may also involve 22 nucleotide-long (22-nt) miRNAs that are able to trigger the conversion of their mRNA targets into double-stranded RNA (dsRNA) molecules, through the recruitment and ensuing action of RNA-dependent RNA polymerase 6 (RDR6). These dsRNAs are then processed by DICER-LIKE (DCL) 4 to generate secondary 21 nt siRNA, which are often phased with respect to the binding site of the miRNA and are known as phasiRNA [24–26]. These secondary phasiRNAs amplify silencing of the target mRNA and may act *in trans* to silence additional mRNAs, not primarily recognized by the miRNA triggering the process. Thus, 22 nt miRNAs can establish regulatory networks/cascades suppressing the expression of several genes by the combined action of primary miRNA and secondary sRNAs. The best-characterized endogenous secondary siRNAs are known as *trans*-acting RNAs (tasiRNAs) and their biogenesis is triggered by 22 nt miRNA-directed cleavage of a non-coding TAS primary transcript[27–32].

Several 22 nt miRNAs have been shown to trigger production of phasiRNAs from target NLR mRNAs in several plant families, *e.g*. Brassicaceae, Coniferae, Fabaceae, Rosaceae, Solanaceae, or Vitaceae (reviewed by [26]). The members of the miR482/2118 families are probably the best characterized and have been identified in several plant species. Several miRNA families, including miR482/2118, trigger phasiRNA production mainly by targeting the NLR gene sequences encoding the P-loop motif, a well-conserved amino acid motif present in the NB-ARC domain of multiple NLRs from *Arabidopsis*, poplar, tobacco and tomato [33–35]. Mutation of a conserved residue within the P-loop results in loss of function of many plant NLRs [3]. MiR482, and related miR472, also target the sequence coding for the P-loop of numerous CNLs and are involved in resistance against bacterial and oomycete pathogens[33–36]. In legumes, several 22 nt miRNAs target NLRs transcripts, mainly CNLs, triggering production of phasiRNAs, leading the authors to hypothesize that miRNAs may act as master regulators of NLR expression via phasiRNA production [23]. In support of this notion, RDR6 has been shown to act as a negative regulator of plant defense against bacterial pathogens in *Arabidopsis* [34].

In this paper, we focus on the molecular characterization of miRNA825-5p, a miRNA produced from *MIR825* and conserved among the *Brasicaceae* familiy, demonstrating its function as a master regulator of TNLs in *Arabidopsis*. *MIR825* also produces a 21 nt mature miRNA, originally identified in *A. thaliana* [37], shown to depend on DICER LIKE (DCL) ribonuclease 1 (DCL1) activity, and to be down regulated during interaction with a non-pathogenic mutant derivative of *P. syringae* [38]. In this study, we show that miRNA825-5p targets a sequence coding for a highly conserved amino acid motif (TIR2) within the TIR domain of the *Arabidopsis AT5G38850* gene. *AT5G38850* is predicted to encode an uncharacterized TNL, which we have named microRNA-silenced TNL 1 or MIST1, and its expression had been previously been shown to be up regulated during infection with *P. syringae,* in the context of induced systemic resistance (ISR) [39]. The TIR2 conserved target motif immediately precedes the catalytic residue for a recently described nicotinamide adenine dinucleotide (oxidized form, NAD+)-cleaving activity, essential for TNL function [40]. We show that miRNA825-5p generates functional secondary phasiRNAs from *MIST1* transcripts capable of *trans*-acting gene silencing that target the coding sequences of a wide network of previously uncharacterized TNL-encoding genes in *Arabidopsis*. Regulation of *MIST1* affects disease resistance against model bacterial pathogen *Pseudomonas syringae*, since inactivation of the miRNA825-5p, using tandem target mimic (STTM) to prevent primary and secondary regulation of the NLR, enhances *Arabidopsis* resistance against bacteria, while overexpression of this miRNA render plants more susceptible to infection. MiR825-5p is expressed in unchallenged adult leaves and its production is down regulated at transcriptional level in response to PAMPs such as bacterial flagellin but also fungal chitin. These results confirm the importance of miRNA-mediated suppression of NLR disease resistance genes, and reveal the existence of a new regulatory network carried out by miR825-5p and key target gene *MIST1* acting as a hub, which includes many uncharacterized-TNL genes in *Arabidopsis* through targeting of a very conserved functional motif, involved in a new, recently identified enzymatic function, essential for TNL defense signaling.

## Results

### MiR825-5p 22 nt miRNA is a candidate regulator of TNLs

We predicted the targets for both 21 nt miR825 and the 22 nt miRNA generated from opposite arms of the same duplex within the MIR825 transcript, using WMD3 (http://wmd3.weigelworld.org/cgi-bin/webapp.cgi) (**Table S1**). Using default parameters, this analysis rendered only three putative targets for 21 nt miR825, which were annotated as encoding a poly(A)-binding protein (*AT1G71770*), and two ubiquitin carboxyl-terminal hydrolase encoding genes (*AT3G47890* and *AT3G47910*) (**Fig. 1A**). These predictions were remarkably more restrictive than previously reported for this miRNA [39]. In contrast, this analysis predicted numerous targets for the 22 nt miRNA derived from *MIRNA825* (top four shown in **Fig. 1A**), and these displayed notably lower hybridization energies and better pairing, and were consistently annotated as uncharacterized TNLs (**Table S1**). Three of these predicted targets have been previously reported to display changes in expression in plants with simultaneously altered levels of both 21 nt and 22 nt miR825s [39]. Target prediction carried out using alternative software (psRNATarget, [41]) also often used in the literature for this purpose rendered little differences (**Table S1**). Predicted processing sites for all putative target TNL genes mapped to the sequence coding the TIR domains, and in particular to a conserved site, previously named TIR2 motif [42], which immediately precedes the catalytic glutamic acid residue for the NAD+-cleavage enzymatic activity recently demonstrated for TIR domains (**Fig. 1B**; [40]). Since the majority of NLRs regulated by 22 nt miRNAs described to date belong to the CNL family [23, 33], and only CNLs have been demonstrated as directly targeted by these miRNAs in *Arabidopsis* [34], this finding further attracted our attention.

**Figure 1.**
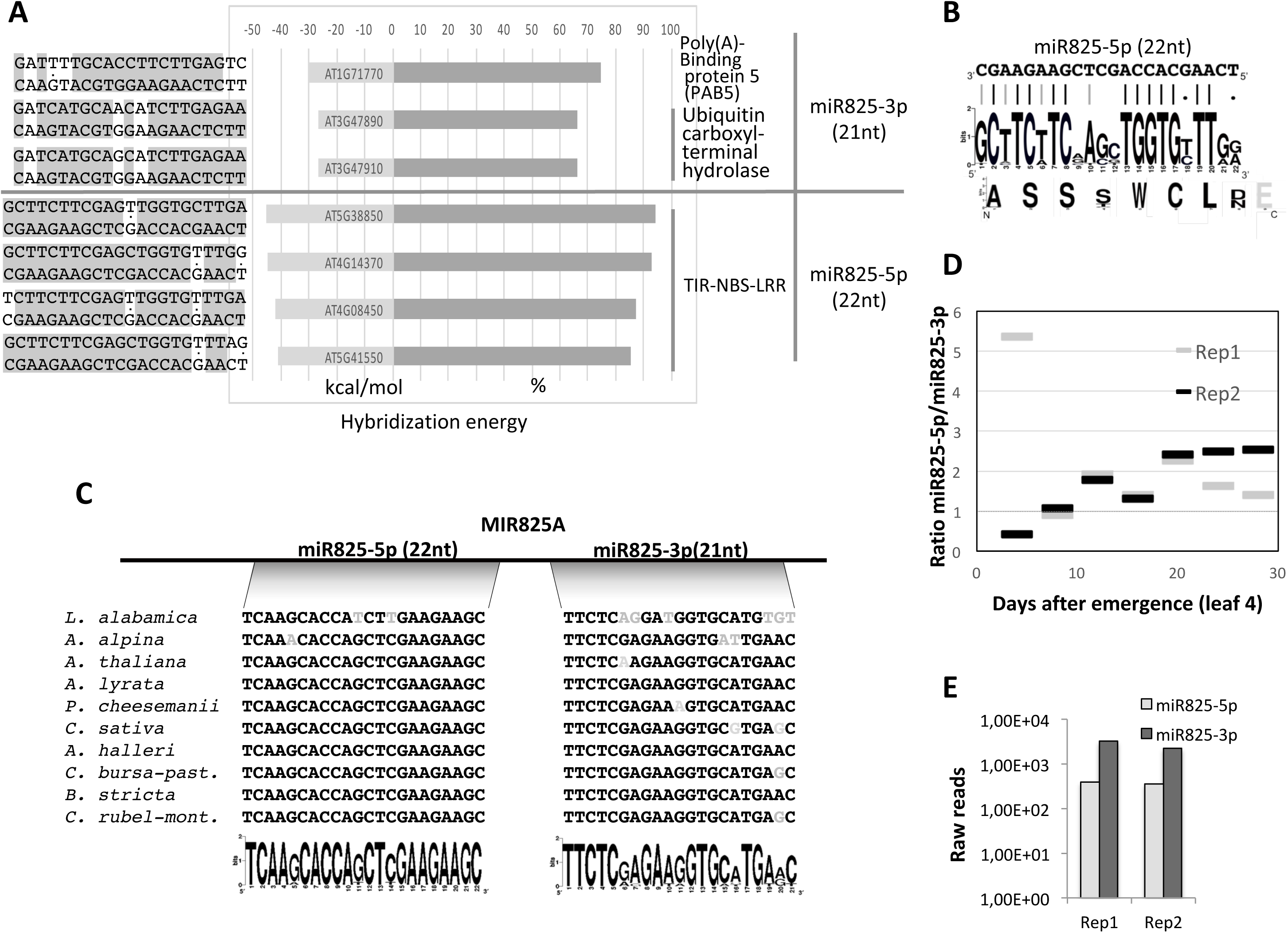
MiR825-5p is a candidate regulator of TNLs. **A** We used WMD3 (Ossowski et al., 2008) and default parameters on Araport11 to predict targets for MIR825-encoded 21 nt (miR825-3p, formerly miR825) and 22 nt (miR825-5p; formerly miR825*). All three predicted targets for 21 nt miR825-3p are shown. Only the top four predicted targets for 22 nt miR825 are shown. Predicted targets for 22 nt miR825 are genes encoding Toll/interleukin-1 (TIR), nuclear binding site (NBS) and leucine-rich repeat (LRR) containing proteins (TIR-NBS-LRR). **B** MiR825-5p sequence paired with the consensus for 18 TNLs (plus one TIR domain protein) putative targets from the *Arabidopsis* genome. The logo corresponding to the consensus protein sequence for miR825-5p target site is shown below (black lines indicates perfect pairing, grey lines perfect pairing with the most conserved nucleotide, and dots indicate variable region that allows pairing at the RNA level). **C** Sequence comparisons between 21 and 22 nt miR825 encoded in different brassica species show that 22 nt miRNA825-5p is conserved to a higher degree than 21 nt miRNA825-3p. **D** Graph shows ratio between levels of miR825-5p (formerly miR825*) and miR825-3p (formerly miR825) in leaf four at different days after emergence in Col-0 plants as stored in public databases (PRJNA186843). Two replicate experiments are shown. Ratios obtained are equal (8 days) or higher than 1.0, except in young leaves (4 days) in one of the two replicates. **E** Graph shows levels of miR825-5p and miR825-3p pulled down as part of AGO1 complexes. Data has been obtained from GSM2787769 and GSM2787770 public databases.

Sequence analysis shows that *MIRNA825* is present mainly in brassica species, with the 22 nt miR825 displaying very high conservation (**Fig. 1C**; [43]). Rather surprisingly a sRNA displaying very high sequence similarity to 22 nt miR825 has been recently reported in *Triticum aestivum L.* [44]. We carried out data mining of public databases including 14 libraries from different developmental stages of *Arabidopsis* adult leaves (Bioproject PRJNA186843), and found that the 22 nt miR825 consistently accumulates to similar or significantly higher levels than the 21 nt generated from the opposite arm of the same duplex (**Fig. 1D**). Interestingly, the *MIRNA825* pair has been previously classified as a Class V miRNA, according to the thermostability of the duplex strands [45]. Class V duplexes differ from the rest in that they form symmetrical miRNA duplexes with equivalent thermostability at the terminus of both duplex strands, resulting in equal accumulation of both miRNA and passenger miRNA. Data mining of public databases showed that 22 nt miR825 is consistently pulled down in association to AGO1 complexes (**Fig. 1E**) supporting the notion of its involvement in gene silencing of its target genes.

In summary, all these data supports the notion of processing of opposite arms of the *MIRNA825* duplex leading to the accumulation of two 21 nt and 22 nt functional miRNAs, thus making the current designation of the 22 nt form as miRNA* or passenger miRNA outdated. Thus, in keeping with the designation of these miRNAs in other brassica species (*e.g.* broccoli [46]; *A. lyrata* **miRBase Release 21**), we will name these miRNAs as miR825-5p (22 nt, formerly miR825*) and miR825-3p (21 nt, formerly miR825)

### MiR825-5p is a negative regulator of plant immunity against *P. syringae*

Prior to characterizing the potential role for miR825-5p as a negative TNL regulator in *Arabidopsis*, we verified an actual involvement of *MIRNA825*-derived miRNAs in *Arabidopsis* defense against *P. syringae* (**Fig. 2**). To do so, we engineered transgenic plants to express an artificial miRNAs (amiRs) [47, 48], where the precursor for miR319 is modified to target and silence pri-miR825, as previously reported for other miRNA precursors (**Fig. 2A**) [49]. As expected, these plants displayed reduced levels of pri-miR825, and reduced levels of both mature miR825-5p and miR825-3p, compared to wild type Col-0 plants (**Fig. 2B**). Upon treatment with flg22 flagellin peptide, a major activator of basal defense (PTI) against bacteria, anti825-expressing plants displayed enhanced defense responses, as shown by higher production of reactive oxygen species (ROS) (**Fig. 2C**), and an increased activation of mitogen-activated protein kinases (MPKs) (**Fig. 2D**). In addition, these plants displayed an increased accumulation of pathogenesis-related 1 protein (PR-1) in response to inoculation with *P. syringae* DC3000 expressing avirulence gene AvrRpt2 (**Fig. 2E**). Finally, as conclusive evidence of MIR825 involvement in plant defense against *P. syringae*, MIR825-silenced plants were more resistant to *P. syringae* DC3000 colonization (**Fig. 2F**). These results are in keeping with a previous report linking overexpression and down regulation of levels of MIR825-derived miRNAs to induce systemic resistance (ISR) within the *Bacillus cereous*/ *P. syringae* system [39].

**Figure 2:**
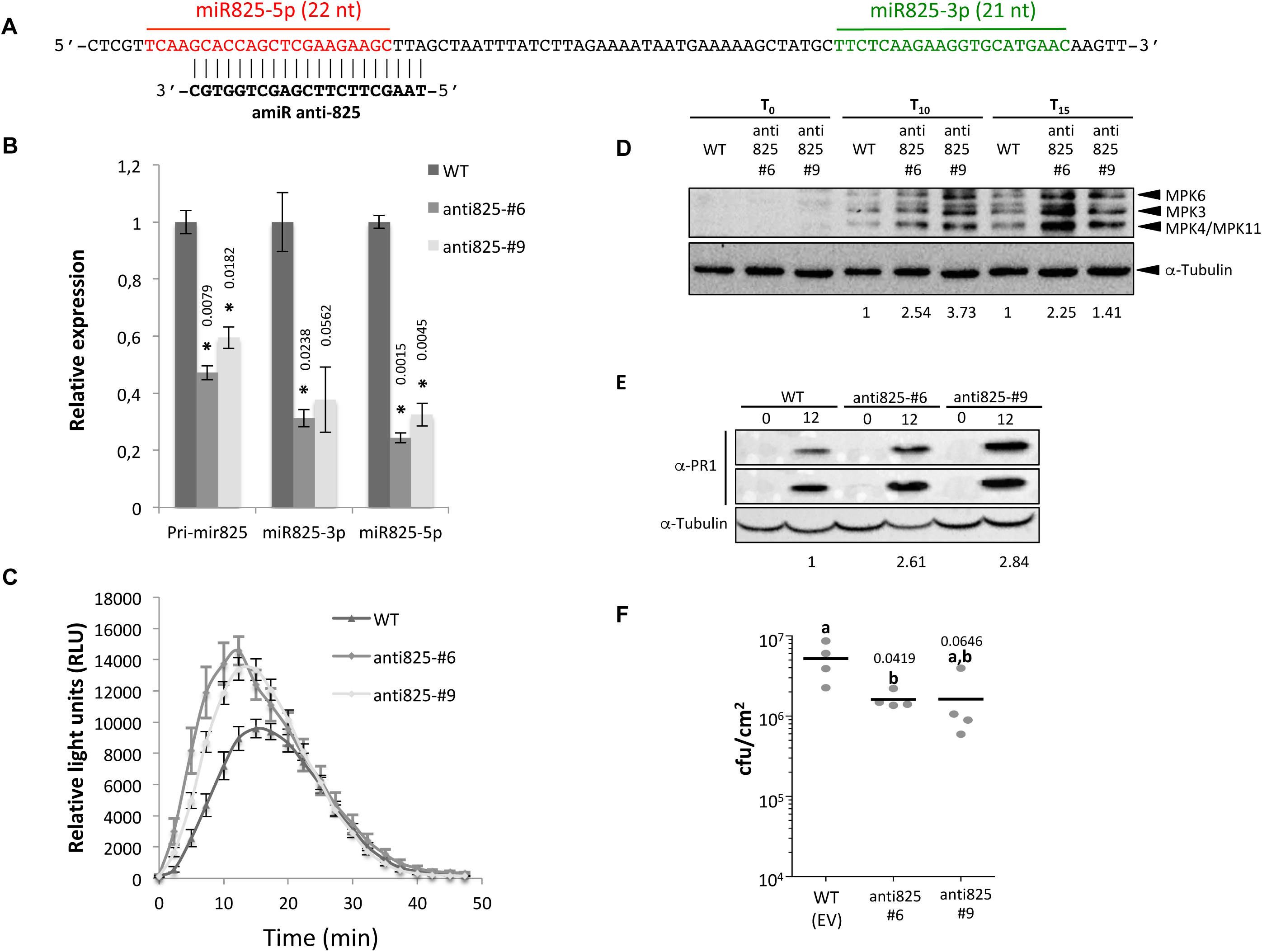
Pri-miR825 has a negative impact on PTI. **A** Sequence of pri-miR825 indicating target for amiR anti825. **B** Plants expressing amiR anti825 display significantly reduced precursor levels, as well as significantly reduced levels of the mature forms of miR825-5p and miR825-3p compared to wild type Col-0 plants (WT). Asterisks indicate results are significantly different from WT plants, as established by a Student’s t-test (P<0.05). Numbers above the bars indicate *P* values. Error bars correspond to standard error. **C** ROS production at different time points after treatment with 100 nM flg22 of WT or anti825 lines. **D** Western Blot analysis showing levels of phosphorylated mitogen-activated protein kinases (MPK3, MPK4, MPK6 and MPK11) after treatment with 100 nM flg22 of wild type (WT) or anti825 lines at three different time points (0, 10 and 15 days post flg22 treatment). Anti-tubulin was used to normalize. Numbers below the blot indicates fold-differences between MPK/tubulin signal ratios calculated using ImageJ (http://imagej.nih.gov/ij/) in anti825 lines and the ratio obtained for WT plants in each time point. **E** Western blot analysis showing PR1 protein levels in WT or anti825 plants after inoculation with 5×10^7^ colony forming units (CFU)/ml of *P. syringae* DC3000 expressing effector AvrRpt2 from a plasmid under a constitutive *nptII* promoter. Samples were taken at 0 or 12 hours post inoculation. Anti-tubulin was used as loading control. **F** Bacterial multiplication assay in WT or antiR825 lines: Leaves were inoculated by infiltration with a solution of 5×10^4^ CFU/ml of *P. syringae* DC3000. Samples were taken 4 days post inoculation and plated. Bacterial counts are shown. Mean values are shown for each plant genotype, although individual values are also represented. Error bars represent standard error. Mean values marked with the same letter were not significantly different from each other as established by Student’s t-test (P<0.05).

Although the large majority of the numerous TNL genes potentially targeted by miR825-5p are uncharacterized, their nature as pathogen receptors, their prevalence among the predicted targets, and the strength of their scores when compared to those of the targets predicted for miR825-3p, suggest that miR825-5p could be responsible for the up regulation of plant immunity against *P. syringae* produced by silencing of pri-miR825 (**Fig. 2**). To test this hypothesis, we generated *A. thaliana* transgenic plants with increased or reduced levels of mature miR825-5p using artificial microRNAs or STTM technology, respectively (**Fig. 3A and B**) [47, 50]. These experimental approaches allow achieving changes in a specific mature miRNA (*i*.*e*. miR825-5p) to be carried out without altering steady state of the additional miRNA generated from the endogenous pri-miRNA duplex (*i*.*e*. miR825-3p) [47, 50]. The transgenic lines displayed significantly increased susceptibility (amiR825-5p; **Fig. 3A**), and resistance (STTM825-5p; **Fig. 3B**) against *P. syringae* DC3000, supporting that miR825-5p acts as a negative regulator of plant immunity against this pathogen. The impact on *P. syringae* colonization of plant leaves of the transgenic lines with reduced levels of mature miR825-5p (**Fig. 3A**) was comparable to that observed in lines with reduced levels of pri-miR825 (**Fig. 2F**), which results in reduced levels of both miR825-5p and miR825-3p. These results support miR825-5p, rather than miR825-3p, as the relevant form of the *MIR825* pair in the regulation of plant immunity against *P. syringae*.

**Figure 3.**
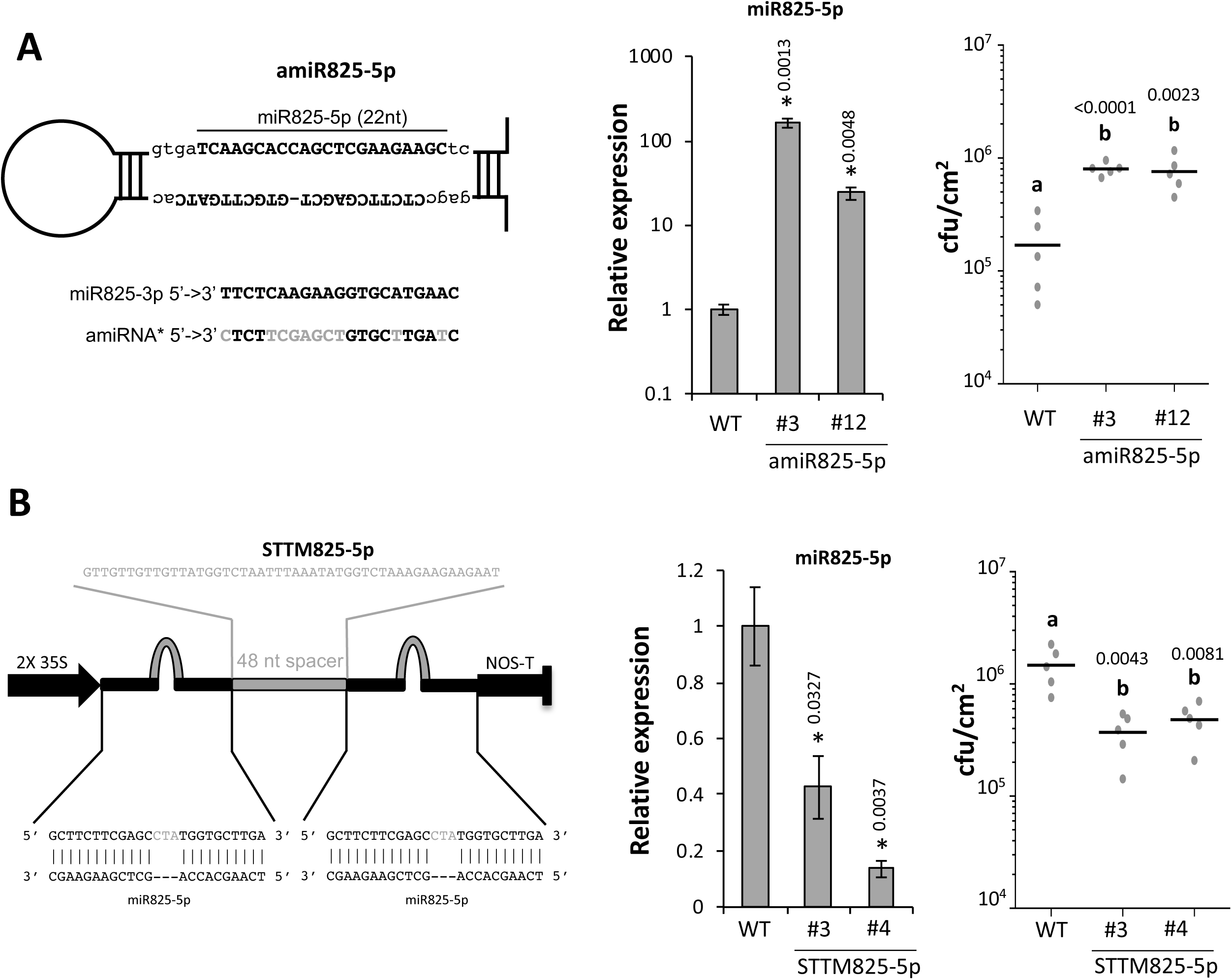
MiR825-5p is a negative regulator of plant immunity against *P. syringae*. **A** Left panel shows sequence and hairpin structure of amiR825-5p. A sequence comparison between miR825-3p and the passenger miRNA generated from this construct (amiRNA*) is shown below the hairpin. Center panel shows relative expression of miR825-5p, while right panel shows bacterial colonization of WT and two independent lines expressing amiR825-5p (#3 and #12). **B** Left panel shows sequence and structure of the STTM825-5p construct. Right panels show relative expression of miR825-5p and bacterial colonization of WT and two independent lines expressing STTM825-5p (#3 and #4). For miR825-5p expression assays, asterisks indicate results are significantly different from WT plants, as established by a Student’s t-test (P<0.05). Error bars correspond to standard error. Numbers above bars indicate the *P* value. For bacterial colonization assays in **A** and **B**, plants were inoculated by infiltration of a 5×10^4^ CFU/ml of *P. syringae* DC3000 solution. Samples were taken 4 days post inoculation and plated. Bacterial counts are shown. Mean values are shown for each plant genotype, although individual values are also represented. Error bars represent standard error. Mean values marked with the same letter were not significantly different from each other as established by Student’s t-test (P<0.05). *P* values are shown above the letters.

### The *AT5G38850* TIR-NBS-LRR transcript is a target of miR825-5p

Our bioinformatics analysis using WMD3 rendered 18 TNL genes, one of which is predicted to produce a truncated version containing only the TIR domain, and another encoding a protein carrying a TIR domain, as potential targets of 22nt miR825-5p (**Fig. 1A; Table S1; Fig. 4A**), with *AT5G38850* as the gene displaying lowest hybridization energy and best pairing. Since some 22 nt miRNAs trigger the production of RNA-derived phased small interfering RNAs (phasiRNAs) from the target sequence [24, 25], and the predicted processing sites for the TNL genes mapped to the conserved TIR domains, we compared the accumulation of sRNAs from each *Arabidopsis* NLRs. We carried out data mining of public databases and found that the number of sRNAs derived from top target *AT5G38850* in adult leaves was the largest for any NLR gene in the *Arabidopsis* genome (**Fig. 4B**). A close examination of sRNAs generated from those that accumulate the most is shown in **Fig. S1**.

**Figure 4.**
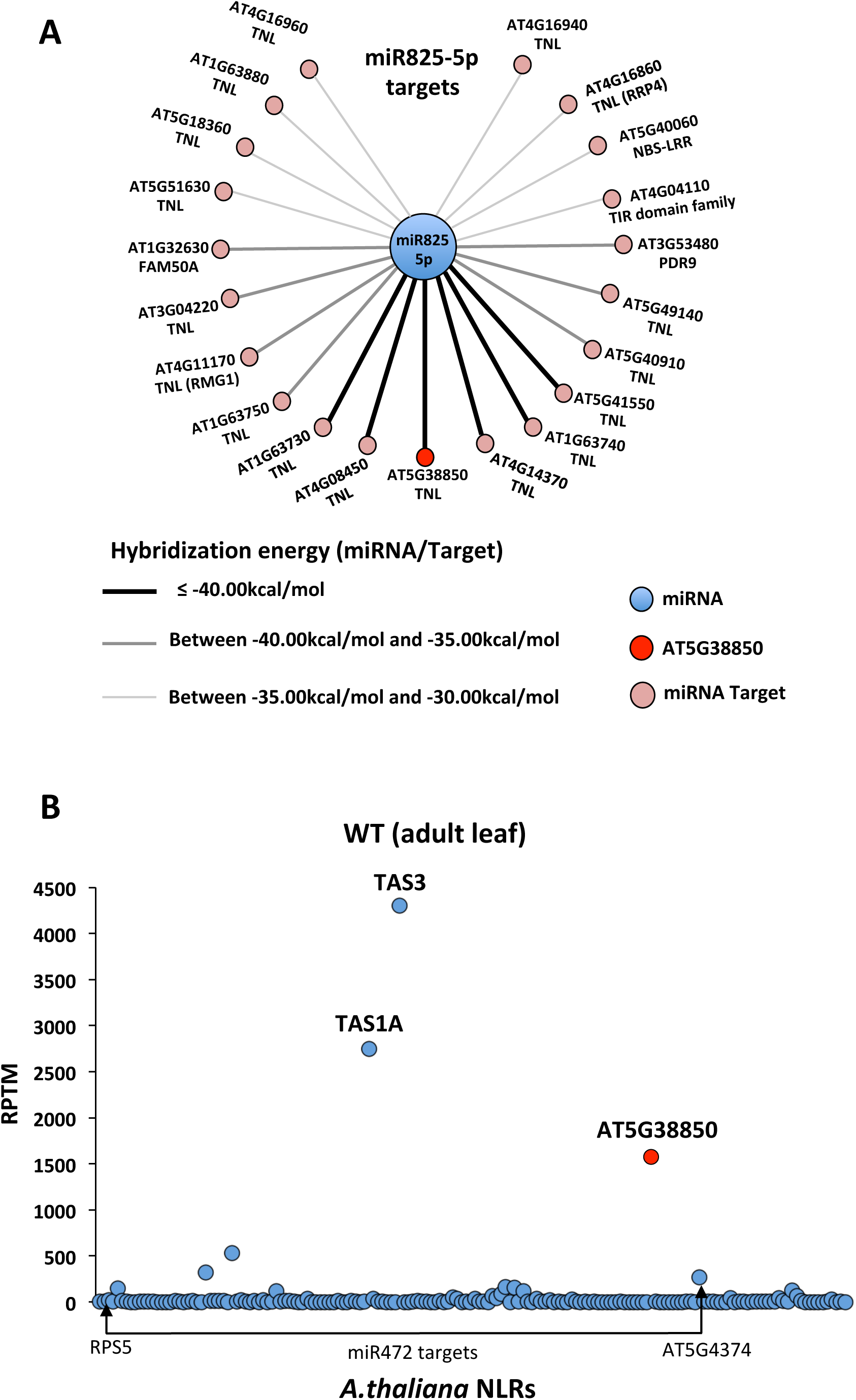
Data mining and regulatory network analysis revealed miR825-5p putative target TNL-encoding *AT5G38850* gene as a central hub for TNL gene regulation. **A** Regulatory network showing all 21 putative targets for miR825-5p as predicted using WMD3 and default parameters on Araport11. TNL is indicated for the 17 out of these 21 that are annotated as such. Two additional genes encode a truncated TIR-NBS-LRR and a TIR domain-carrying protein. **B** Graph shows sRNA accumulation from all NLRs within the *Arabidopsis* genome (data obtained from NCBI: BioProject SRP097592, WT library). Graph displays number of sRNA (reads per 10 million small RNAs mapped, RTMP) accumulated from each NLR-encoding gene. Number of sRNA accumulated from *TAS1A* and *TAS3* are provided as reference.

Processing of the miR825 duplex has been proposed to depend on the activities of DCL1 and DCL3 [38, 51]. Thus, as a first step in the characterization of miR825-5p targets, we confirmed that accumulation of pri-miR825 increased in a *dcl1-7* mutant (**Fig. 5A**), while levels of miR825-5p were significantly reduced. As expected from the notion of miR825-5p targeting *AT5G38850*, transcript levels of this gene were significantly increased. To seek direct validation of miR825-5p regulation of *AT5G38850*, we generated a gene fusion of the genomic sequence of the *AT5G38850* (including its own 5’-UTR region, exons and introns) to the Green Fluorescent Protein gene (*GFP*) ORF, to be transcribed under the control of a 35S constitutive promoter (wt-*AT5G38850*; **Fig. 5B**). As control, we generated a modified version of this gene fusion in which the miR825-5p target site is no longer complementary to this miRNA without altering the corresponding amino acid sequence (m-*AT5G38850*; **Fig. 5B**). *N. benthamiana* leaves co-expressing wt-*AT5G38850* and miR825-5p accumulate very low levels of GFP, when compared to those accumulated in leaves co-expressing the gene and unrelated miR319 (**Fig. 5C**). Furthermore, GFP levels of control leaves expressing wt-*AT5G38850* and miR319 were similar to those detected in leaves co-expressing m-*AT5G38850* and either miR825-5p or miR319 (**Fig. 5C**). These results indicate that miR825-5p targets and regulates mRNA accumulation of *AT5G38850* by recognizing a complementary sequence located at the TIR domain. In support of this notion, we measured the level of expression of endogenous *AT5G38850* in relation to levels of miR825-5p in previously generated transgenic plants displaying altered levels of this mature miRNA (**Fig. 3**). In keeping with results shown in **Fig. 5C**, accumulation of endogenous *AT5G38850* transcripts displayed a negative correlation with levels of miR825-5p in all different genotypes tested (**Fig. 5D**). Based on these results, we named *AT5G38850 MIST1* for <u>mi</u>RNA-<u>s</u>ilenced <u>T</u>NL-1, and will refer to it as such hereafter.

**Figure 5.**
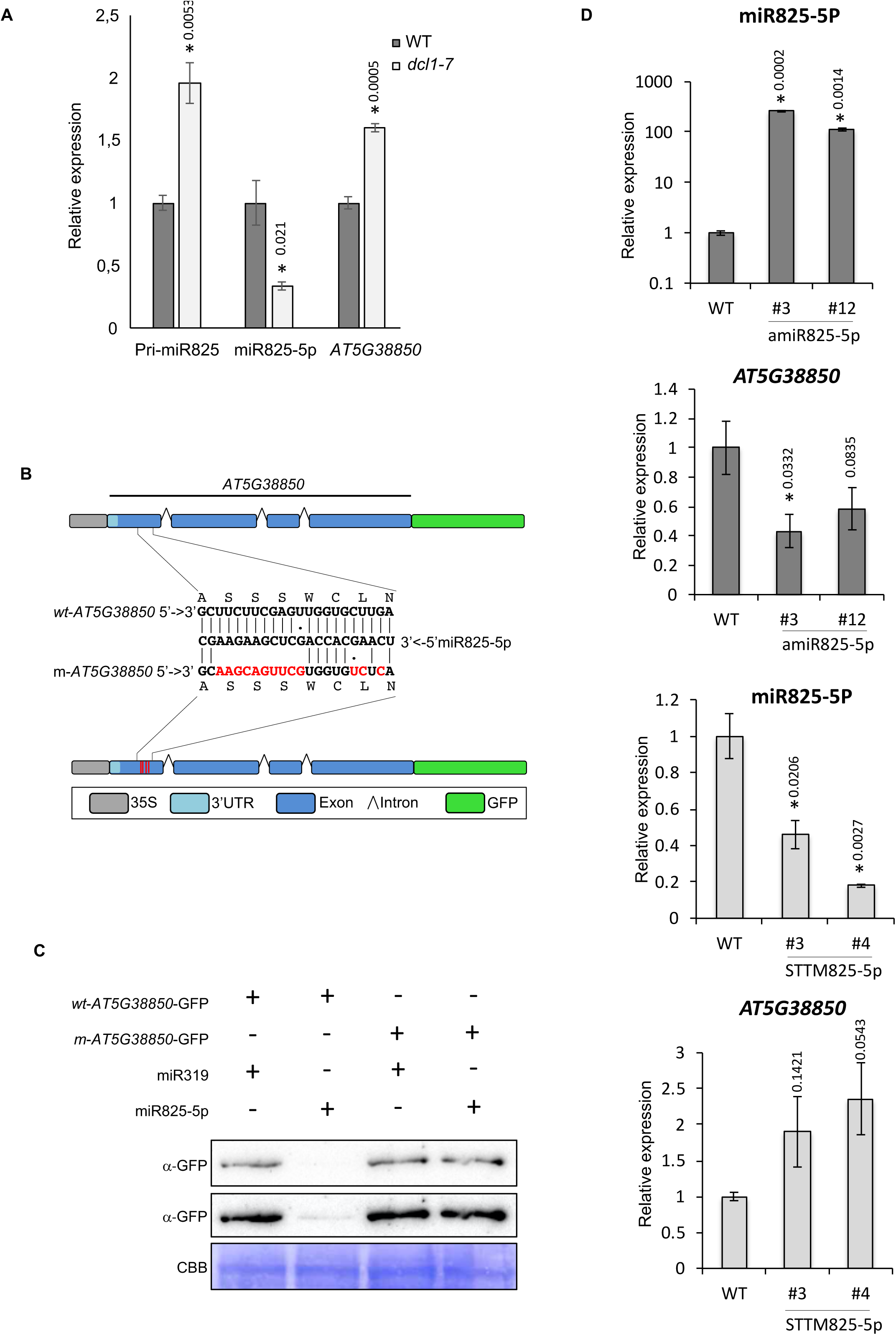
TNL-encoding *AT5G38850* gene is a target for miR825-5p regulation. **A** Graph shows levels of pri-miR825, miR825-5p and *AT5G38850* mRNA in WT *versus dcl1-7* mutant plants. Asterisks indicate results are significantly different from WT plants, as established by a Student’s t-test (P<0.05). Error bars correspond to standard error. **B** Gene fusion of *AT5G38850* (includes its own 3’-UTR region, exons and introns) to the Green Fluorescent Protein gene (*GFP*) ORF. The gene fusion is under the transcriptional control of a 35S constitutive promoter (wt-*AT5G38850*). Modified version carries mutations making the transcript generated no longer complementary to miR825-5p, without affecting protein coding (m-*AT5G38850*). **C** Western Blot analysis using an anti-GFP antibody of *Nicotiana benthamiana* leaves transiently co-expressing either wt-*AT5G38850* or mut-*AT5G38850,* with either miR825-5p or unrelated miR319. Coomassie blue staining of the membrane is shown as loading control. **D** Accumulation of endogenous *AT5G38850* transcripts negatively correlated with levels of miR825-5p in all different genotypes tested. Asterisks indicate results are significantly different from WT plants, as established by a Student’s t-test (P<0.05). Error bars correspond to standard error.

### MiR825-5p triggers phasiRNAs production from target *MIST1* transcripts

Data mining results show that sRNA accumulation derived from *MIST1* is notably larger than those derived from any of the other NLR-encoding candidate targets genes of miR825-5p (**Fig. 4B**), including the known CNL-encoding gene targets of the RDR6-mediated regulation triggered by miR472 (*RPS5*, *RSG1* and *AT5G43740*) [25,34,52]. MiR825-5p is predicted to arise from an asymmetric fold-back precursor containing asymmetric bulges (**Fig. 6A and B**), a characteristic associated to the ability to 22 nt miRNA to trigger phasiRNA production from complementary targets [24,25,53], thus suggesting miR825-5p as a potential trigger of phasiRNA production from *MIST1*, in keeping with previously reported computational predictions [24, 54]. Interestingly, a previous study found a positive correlation between levels of this miRNA and accumulation of siRNAs from *MIST1* during induced systemic resistance (ISR) [39], however changes in RDR6, such as those reported during PTI [34], could be responsible for this correlation. Close examination of the sRNAs produced from *MIST1* supports the role of miR825-5p as a trigger of phasiRNA on this locus: (*i*) the first sRNA that accumulates at significant levels from the positive strand of *MIST1* maps between positions 10-11 after the predicted cleavage site of miR825-5p in *MIST1* (**Fig. 6C, D and E**), and (*ii*) sRNAs that accumulate from this cleavage site onwards displays regular spacing befitting DCL4-mediated production of phased RNAs (**Fig. 6F**). Interestingly, Branscheid and collaborators [55] reported that siRNAs are produced from the target fragment that displays the least stable base pairing to the miRNA. In the case of miR825-5p pairing with *MIST1* mRNA, this would be the 3’ target fragment (**Fig. 6G**), in keeping with miR825-5p production of sRNAs from *MIST1* taking place at the 3’ side of the miR825-5p target sequence.

**Figure 6.**
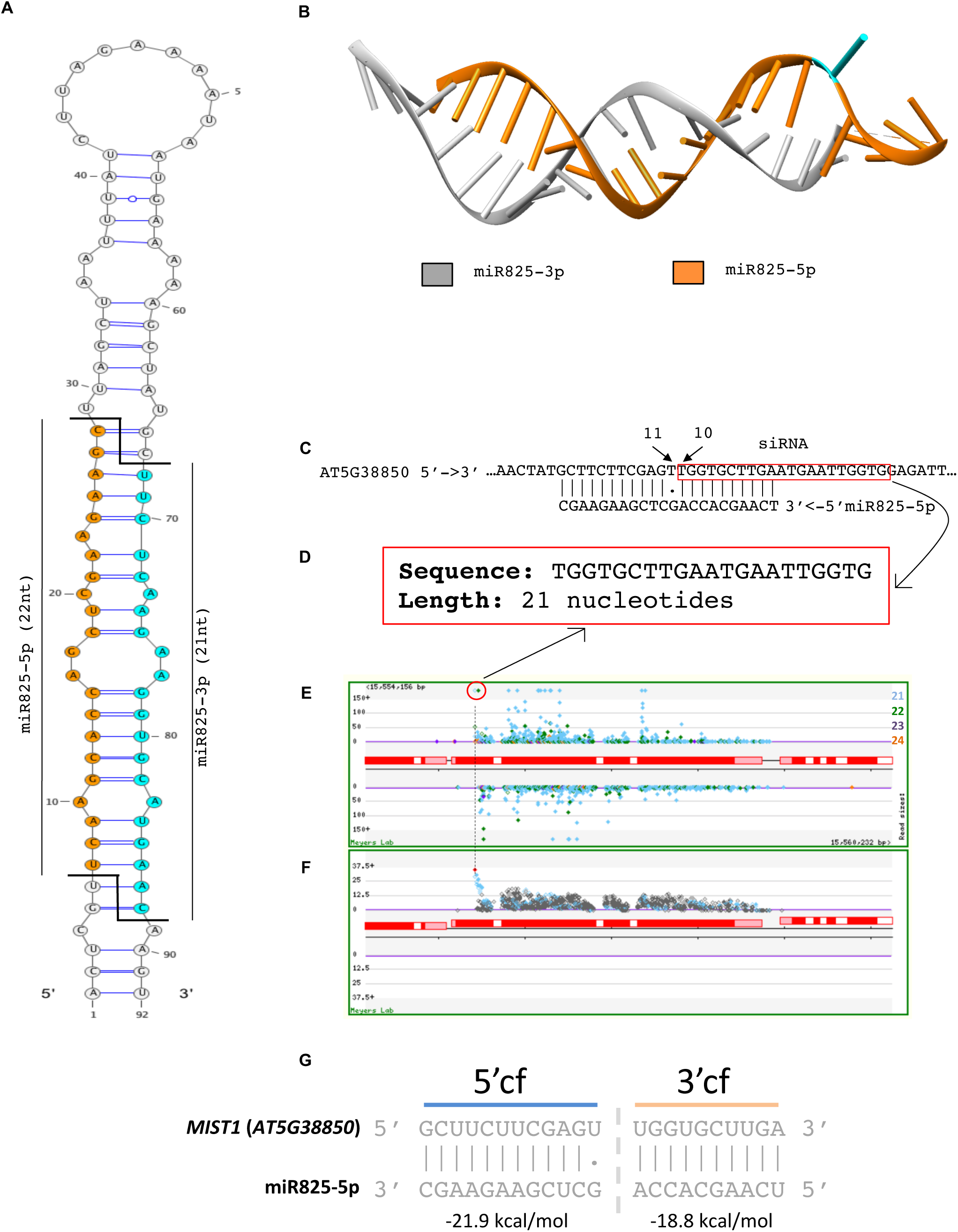
MiRNA825-5p is a putative trigger for phasiRNAs production from *MIST1* transcripts. **A** Pri-microRNA825 predicted hairpin structure. Secondary structure was predicted using Mfold and visualized with Varna. MiR825-5p sequence is indicated in orange. MiR825-3p sequence is indicated in blue. **B** Predicted 3D-structure for the asymmetric duplex formed between miR825-5p and miR825-5p. Predictions were done using RNAfold and MC-fold/MC-Sym pipeline. **C** Sequence complementarity between miR825-5p and its target site in *MIST1*. Sequence matching the first sRNA that accumulates from this transcript is highlighted. Its position matches that predicted for the first phasiRNA to be generated after cleavage by RISC-miR825-5p, between nucleotides 10 and 11 nucleotides. **D** Sequence and length of the sRNA highlighted in **C**. **E** Screenshot from MPSS showing sRNAs that accumulate from *MIST1* (*AT5G38850)*. **F** Screenshot from MPSS showing the Phasing Analysis for the region analyzed in **E**. **G** Predicted free energy for hybridization between miR825-5p and 3’ target fragment/ 5’ target fragment of *MIST1*. Prediction was done using UNAFold as previously described by Branscheid and collaborators (**2015**).

Additional evidence to support that siRNAs from *MIST1* are the product of a canonical biogenesis pathway of tasiRNAs/phasiRNAs (RDR6-DCL4 dependent) can be gathered from the accumulation of *MIST1*-derived sRNAs in adult leaves as described in public libraries for *dcl2/4* and *rdr6 Arabidopsis* mutants (SRA Bioproject: SRP097592). Data mining of siRNA accumulation in single and double mutants revealed that accumulation of 21/22 nt siRNAs derived from both plus and minus strand of *MIST1* require the function of RDR6 (**Fig. 7A and B**), as previously reported for tasi/phasiRNAs production [27–30].

**Figure 7.**
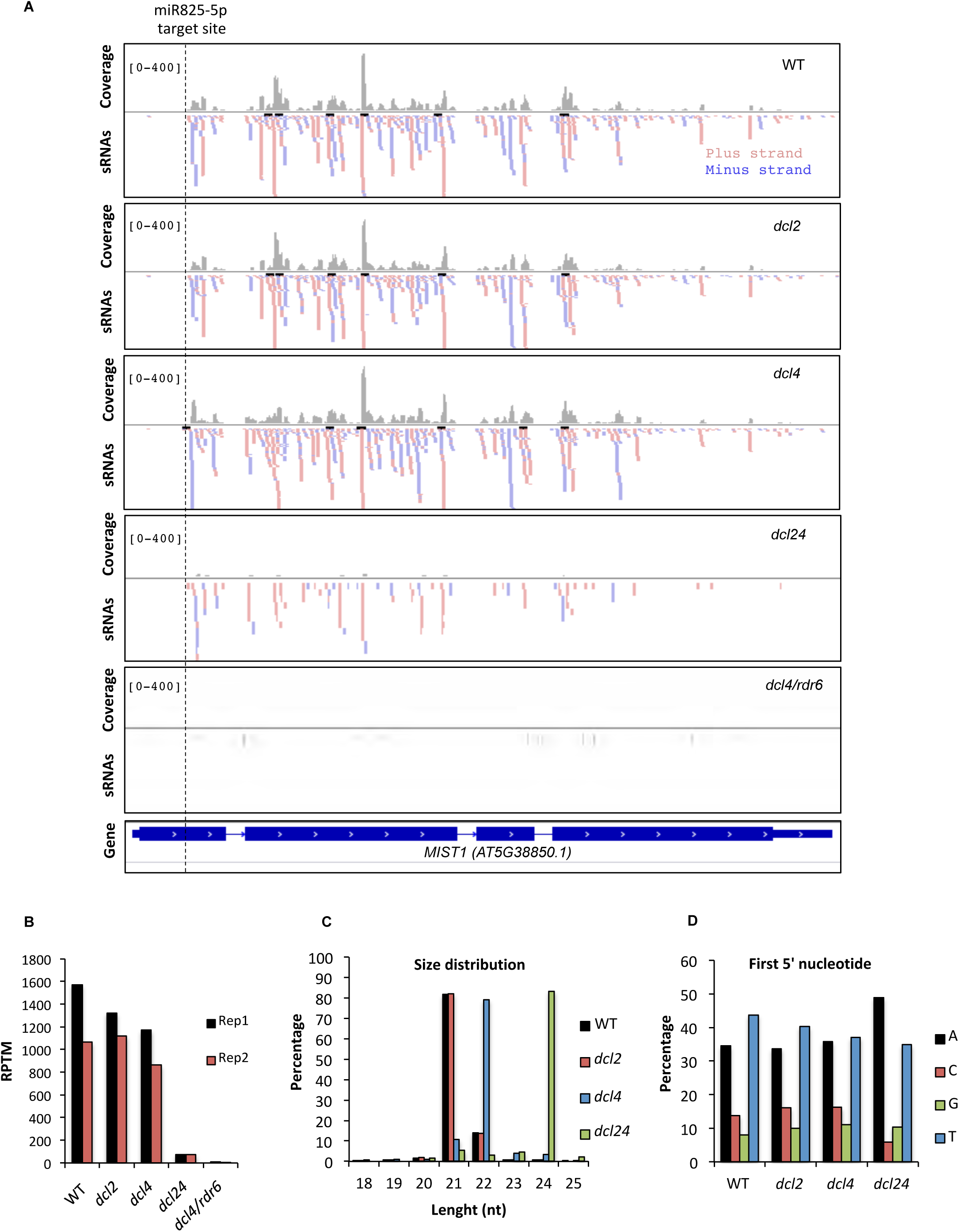
Distribution and DCL/ RDR6-dependency of *MIST1-*derived siRNAs. **A** IGV screenshots showing sRNAs production from *AT5G38850* genomic region, in wild type plants (WT), *DCL2* mutant (*dcl2*), *DCL4* mutant (*dcl4*), *DCL24* double mutant (*dcl24*) and *DCL4/RDR6* double mutant (*dcl4/rdr6*) plants. The dashed line represents miR825-5p target site. Coverage is indicated in grey, and red and blue are used to indicate sRNAs generated from either the positive or negative strand, respectively. The gene model is represented at the bottom. **B** Quantification of sRNAs produced from *MIST1* as RPTM (Reads Per Ten Millions) in two independent biological replicates (replicate 1 represented in **A**). **C** Size distribution (as percentage) of sRNAs that map to *MIST1*. **D** Analysis of first 5’ nucleotide (as percentage) present in the sRNAs that map to *MIST1*. In **C** and **D** results from the *dcl4/rdr6* mutant are not represented due to total absence of sRNAs accumulating from this region.

Mutation of either *dcl2* or *dcl4* causes a small decrease in siRNA levels when compared to the wild type (on average 93% and 77% of wild type levels, respectively), but the combination of the *dcl2* and *dcl4* mutations causes a dramatic reduction on the accumulation of siRNA from this locus (on average less than 6% of wild type levels) (**Fig 7B**). Indeed, siRNAs accumulating in the *dcl2/4* double mutant is limited to DCL3-derived 24 nt siRNAs (**Fig. 7C**) supporting a degree of functional overlap between DCL2 and DCL4 consistent with previously published reports [56, 57]. Percentages of first nucleotide composition were similar in wild type, *dcl2* and *dcl4* single mutants and different from those detected in the *dcl2/4* double mutant (**Fig. 7D**). The abundance of Ts and As in the first nucleotide of the wild type populations suggests these sRNAs could be potentially loaded onto AGO1 and AGO2, respectively[58].

Finally, we used amiR825-5p and STTM825-5p lines, which display elevated and decreased levels of mature of miR825-5p, respectively (described in **Fig. 3A**), to analyze production of phasiRNAs from *MIST1* (**Fig. 8**). Using small RNA Northern blot analysis, we found that accumulation of phasiRNAs from this locus (**Fig. 8A**) was directly correlated with the level of expression of mature miR825-5p in each of the lines tested: a faint band could be seen in wild type plants, and even fainter bands in STTM825-5p plants, while stronger bands were detectable in the amiR825-5p lines, in correspondence to their respective levels of expression (**Fig. 3A and Fig. 5D**). Furthermore, phasiRNAs derived from *MIST1* are found to be loaded onto AGO1 and AGO2 complexes in basal conditions in wild type plants (**Fig. 8B, C and D**), supporting their potential involvement in transitive silencing.

**Figure 8.**
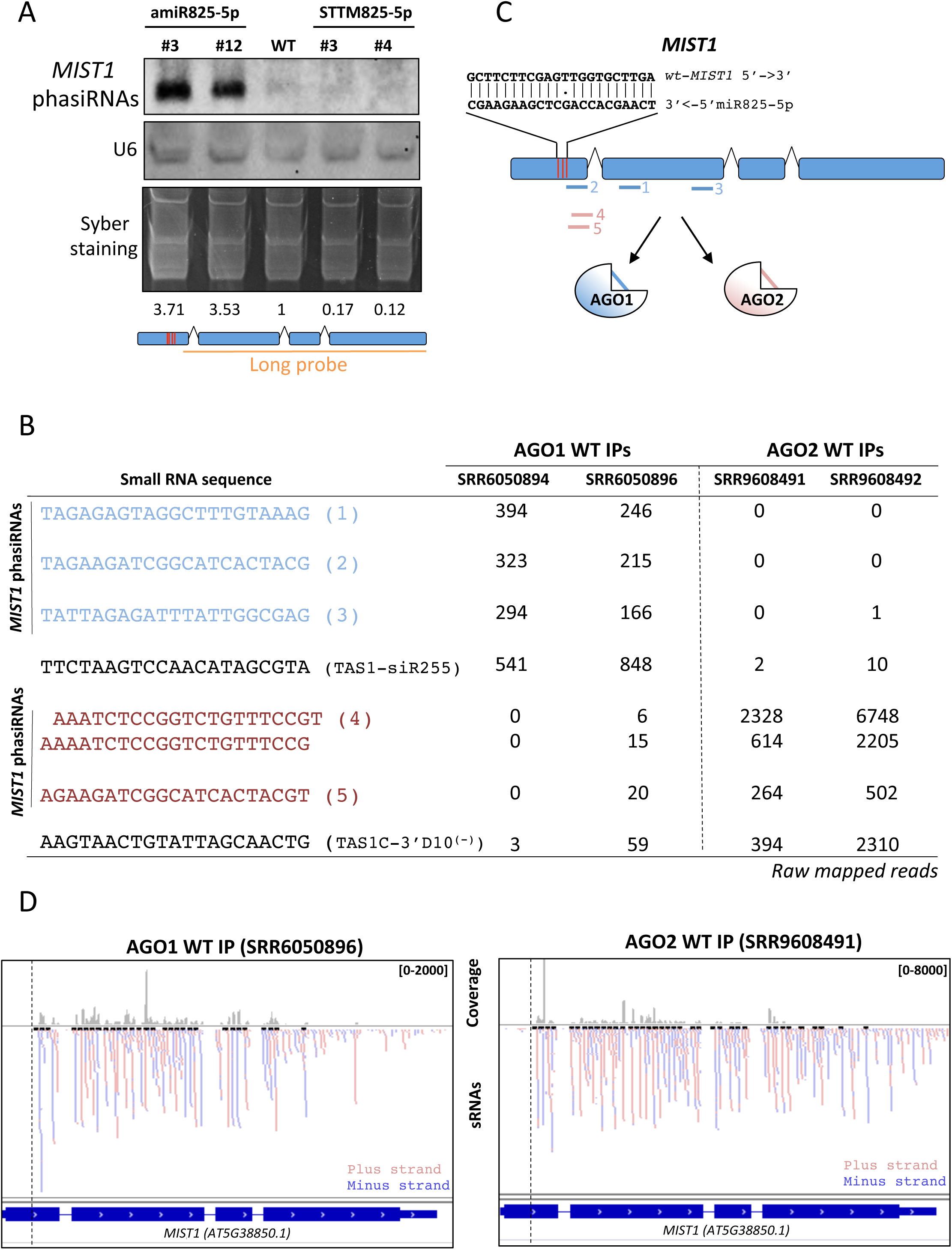
*MIST1*-derived phasiRNAs accumulate according to miR825-5p levels and are loaded onto AGO1 or AGO2 complexes. **A** SiRNA accumulation from *MIST1* shown by small RNA Northern blot analysis. Probe used is displayed. U6 and Syber staining were used to normalize **B** Table shows raw mapped reads for *MIST1*-derived most abundant phasiRNAs in AGO1/AGO2 pull down experiments. Corresponding libraries are indicated. Well-established secondary tasiRNAs generated from *TAS1* genes are included as references. **C** Localization within MIST1 of phasiRNAs included in B as pulled down with AGO1 or AGO2 complexes. MiR825-5p target site is included as a reference. **D** IGV screenshot showing sRNAs production from *MIST1* genomic region that are pulled down with AGO1 and AGO2 complexes. Corresponding libraries are indicated.

### MiR825-5p-triggered phasiRNAs produced from *MIST1* can act *in trans* to silence gene expression

To confirm that miR825-5p-triggers generation of phasiRNAs from its *MIST1* target site, we used miRNA-Induced Gene Silencing technology (MIGS) [59] to generate *Arabidopsis* transgenic plants expressing a gene fusion between the miR825-5p *MIST1* target sequence and *Arabidopsis AGAMOUS* gene under a CaMV 35S promoter (MIGSS825-5pTS; **Fig. 9A**). Thus, if miR825-5p triggers siRNA production at this site, it would lead to the generation of phasiRNAs from the *AGAMOUS* transcript that would silence *AGAMOUS* expression and cause typical flower phenotypes associated to lack of *AGAMOUS* function [59, 60], thus acting as a proxy to demonstrate miR825-5p phasiRNA-mediated transitivity. Transgenic plants expressing MIGSS825-5pTS displayed no apparent flower phenotype (**Fig. 9B and C**). This result could be due to levels of mature miR825-5p in flowers not being sufficient to silence the highly expressed *AGAMOUS* gene [61]. Levels of mature miR825-3p have been reported to be significantly lower in flowers than in leaves [51]. To cover this eventuality, we crossed our lines carrying the *AGAMOUS* sensor system with those expressing miR825-5p from the CaMV 35S within the amiR825-5p construct described above (**Fig. 3A**). These plants, carrying both constructs (MIGSS825-5pTS and amiR825-5p) displayed flower phenotypes typically caused by a mild to moderate silencing of the *AGAMOUS* gene [62], including distorted pistil, lack of maturation of the stamens, and infertility (**Fig. 9B and C**). Control plants expressing the amiR825-5p construct only displayed wild type flower phenotypes (**Fig. 9B and C**). This is in keeping with miR825-5p triggering *in trans* silencing of endogenous *AGAMOUS* when acting at the target site of *MIST1*.

**Figure 9.**
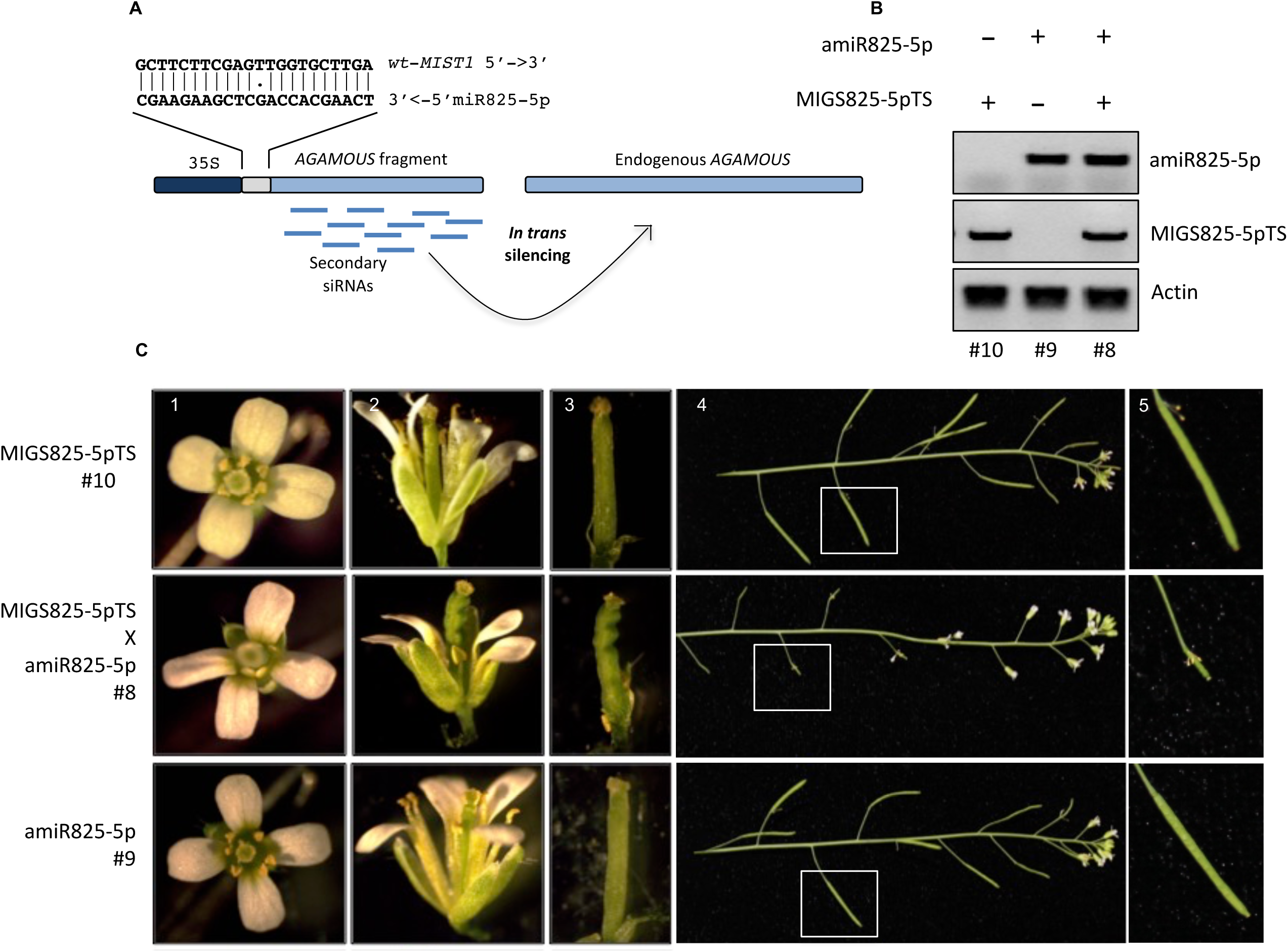
MiR825-5p triggers transitivity at *MIST1* target site. **A** Experimental design for MIGSS825-5p-TS. MiR825-5p target site from *MIST1* is fused to a 500 bp fragment of *AGAMOUS* gene and expression of the construct driven by a 35S constitutive promoter. Recognition by the RISC-miR825-5p complex is expected to triggers phasiRNAs production from the AGAMOUS fragment, which are expected to to silence the endogenous *AGAMOUS* gene *in trans*. **B** DNA genotyping of plants with the amiR825-5p (#9), MIGS825-5pTS (#10). **C.** Flower phenotypes for the different genotypes.

### The *MIR825* promoter is down regulated in response to perception of pathogen associated molecular patterns (PAMPs)

Down regulation of plant immunity against *P. syringae* by miR825-5p and phasiRNAs generated from *MIST1* could be lifted during pathogen interaction by different mechanisms. Since levels of mature miR825-3p, whose targets are not seemingly linked to immunity, have been previously reported to decrease following bacterial perception [38,39,63] we investigated whether *MIR825* could be regulated at a transcriptional level. Inoculation with *P. syringae* DC3000 caused a 70% decrease in pri-miR825 accumulation at 3 hours post-inoculation (**Fig. 10A**). A similar reduction was observed after treatment with the elicitor flagellin peptide flg22 (**Fig. 10B**), suggesting flagellin perception is responsible for the down regulation observed following bacterial entry. However, this down regulation is not specific to flagellin perception since a similar decrease in the accumulation of pri-miR825 is observed in Col-0 *fls2* mutant plants (unable to perceive flagellin) with *P. syringae* DC3000 (**Fig. 10C**), indicating that regulation of pri-miR825 levels is part of PAMP-triggered immunity, and others PAMPs present in this pathogen can trigger a similar effect. To determine if pri-miR825 down regulation was specific to bacterial PAMP-perception, we treated plants with chitin, a fungal PAMP equivalent to bacterial flagellin (**Fig. 10D**), and found that pri-miR825 was similarly reduced after treatment, supporting that pri-miR825 expression responds to general PAMP perception. To follow the dynamic of changes on pri-miR825 levels following PAMP perception, we carried out a time-course experiment upon flg22 treatment using RT-*q*PCR (**Fig. 10E**). The results showed a drastic drop in pri-miR825 accumulation at 3h after treatment, followed by a slow progressive recovery, reaching levels similar to those detected before PAMP perception 48h later.

**Figure 10.**
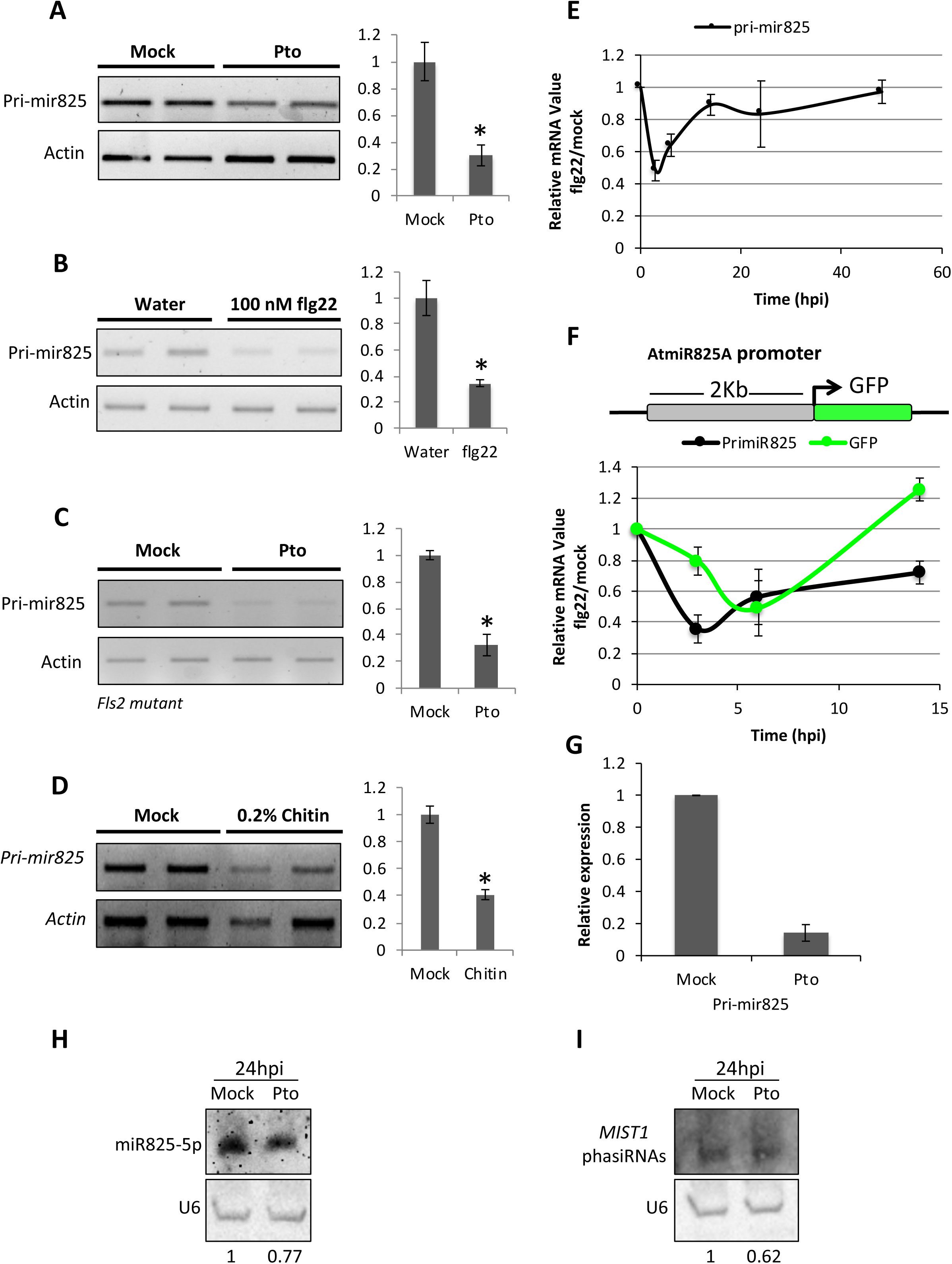
Pri-miR825 is transcriptionally down regulated upon PAMP-perception. **A-D** Semi quantitative RT-PCR show levels of pri-miR825 3 hours: **A** post-inoculation with 5×10^7^ CFU/ml of *P. syringae* DC3000. **B** post-treatment with flg22, **C** post-inoculation of *fls2* mutant plants with, 5×10^7^ CFU/ml of *P. syringae* DC3000, and **D** post-treatment with chitin. Accompanying graphs correspond to Image J quantification of the bands. **E** Time course experiment using RT-*q*PCR to follow accumulation of pri-miR825 transcripts after flg22 treatment. **F** Time course experiment using RT-*q*PCR to follow accumulation of endogenous pri-miR825 transcripts and that of transcripts from the transgene formed by fusión of the AtMIR825 promoter to the GFP coding sequence, after flg22 treatment in a pMIR825*::GFP* transgenic line. **G** Northern blot analysis of the levels of miR825-5p in sRNA samples taken from adult leaves 24 hours post-inoculation with either Pto DC3000 (Pto) or the inoculating solution (mock). **I** Northern blot analysis of the levels of sRNAs producted from the *MIST1* transcript in sRNA samples taken from adult leaves 24 hours post-inoculation with either Pto DC3000 (Pto) or the inoculating solution (mock).

Down regulation of miRNA precursor levels can be established at different levels. To determine if transcriptional down regulation of the *MIR825* gene promoter is involved in down regulation of pri-miR825 levels upon PAMP-perception, we generated *A. thaliana* transgenic lines expressing the *GFP* gene under the control of the *MIR825* promoter (**Fig. 10F**). Western-blot analysis of GFP levels in several independent transgenic lines, confirmed GFP protein accumulation in adult leaves (**Fig. S1**). Treatment of these lines with flg22 determined similar dynamics between GFP mRNA and pri-miR825 transcript accumulation (**Fig. 10F**). A slight delay of GFP mRNA level accumulation is observed, perhaps due to the processing of pri-miR825 by DCL1, a process likely to contribute to a quicker reduction of the precursor. Interestingly, although levels of pri-miR825 had recovered considerably 24 h after treatment with flagellin, these are still significantly lower 24 hours post-inoculation (hpi) with *P. syringae* DC3000 (**Fig. 10G**). Levels of miR825-5p also displayed a decrease 24 hpi, although this decrease was not as strong as that seen for pri-miR825 levels (**Fig. 10H**). A similar trend could be observed for phasiRNAs generated from *MIST1* transcripts at both time points (**Fig. 10I**).

## Discussion

The current theory on the evolution and maintenance of stable resistance polymorphism requires cost of resistance and/or virulence acting in combination with frequency-dependent selection [18, 64]. Pathogen resistance can have two distinct types of fitness costs: (*i*) the cost of surveillance, which results from harboring R genes in anticipation of pathogen attack [65], and (*ii*) the cost of defense that accrues from the activation of resistance during the actual pathogen attack [16, 65]. The cost of surveillance is measured in the absence of disease and has been established in *Arabidopsis* for two R genes: *RPS5* and *RPM1* in [15, 17]. MiRNA-mediated regulation of NLRs has been proposed to act mainly by lowering the cost of surveillance, keeping down the expression of regulated R genes, and preventing accidental activation of their expression in the absence of a pathogen threat [34, 35]. Among the miRNA demonstrated to regulate expression of NLRs, 22nt miRNAs seem to be key elements, since they have the potential to establish a wider suppression of the resistance genes by triggering the production of secondary siRNA from primary target genes. The fact that RDR6 is down regulated during PTI and that *rdr6* mutants present stronger ROS and callose deposition upon flagellin treatment and are primed for expression of PTI marker genes [34], strengthens the significance of this mechanism in plant immunity. NLRs have been classified as phasiRNA or phasiNLR-producing loci when 10 or more 21-22 nt phased siRNAs accumulate from its sequence [35]. Whereas numerous CNL genes have been shown to give raise to accumulation of phasiNLRs in many plant species, just a few examples of sRNA accumulation have been reported for TNL genes [23,26,33–36,66].

In this paper, we characterize miR825-5p, a 22 nt miRNA that controls the expression of numerous TNLs genes in *Arabidopsis* through primary targeting of a highly conserved motif (TIR2) within the TIR domain recently linked to a novel enzymatic function essential for defense signaling [40, 42]. The differences in number, scores, pairing and annotations of the predicted targets for each of the two miRNAs derived from *MIR825* (5p/3p), their different degrees of conservation, as well as miR825-5p accumulation in adult leaves and association to AGO1 complexes (**Fig. 1**), convinced us that the initial denomination of 22 nt miR825 as a passenger miRNA (miR825*) was outdated. The fact that the contribution of *MIR825* to plant defense against *P. syringae* (**Fig. 2**) can be recapitulated by miR825-5p alone (**Fig. 3**) reinforced that conclusion.

Our interest in miR825-5p role in regulating immunity was spurred by its characteristics consistent with an ability to trigger RDR6/DCL4-mediated production of phasiRNAs from complementary targets [24,25,53], since it is a 22 nt miRNA predicted to be produced by DCL1 from an asymmetric fold-back precursor containing asymmetric bulges (**Fig. 6A**). Accumulation of sRNA from its potential targets led us to close in top target *MIST1* (*AT5G38850*) (**Fig. 1 and Fig. 5**) as a potential regulatory hub through the production of phasiRNAs (**Fig. 4, and Fig. 6**). Evidence supporting this notion is found in: (*i*) the accumulation of siRNAs from *MIST1* and its pattern in relation to the miR825-5p target site in this locus (**Fig. 6**), (*ii*) its dependency on the function of RDR6 and that of DCL2/DCL4 (**Fig. 7**), and (*iii*) in its correlation to miR825-5p levels and loading into AGO1/AGO2 complexes (**Fig. 8**).

MIST1 displays the domain structure of a canonical TIR-NB-ARC-LRR. The fact that MIST1 contains in its TIR domain the conserved putative catalytic glutamic acid and neighboring residues recently described by [40], as well as a canonical P-loop (Walker A) motif in its NB-ARC domain, suggests that it is an active NLR, and not only a hub for the generation of phasiRNAs. TIR domains are susceptible of self-association, which are required for the essential NADase activity displayed by active plant TNLs [40, 67]. A structural-homology search with MIST1 using the Phyre2 web portal for protein modeling [68] returned several plant TIR domains with high confidence. The best hit corresponds to the TIR domain of the TNL SNC1 [69, 70], predicted with 100% confidence over 155 amino acid residues (8-163), encompassing the MIST1 predicted TIR domain; SNC1 is included amongst the structure-based phylogeny of proteins similar to hSARM1^TIR^ as described by [67]. The available data on TIR-TIR interactions is compatible with their association into high-order oligomers, stabilized in activated NLRs by self-association of other domains such as NB-ARC [71]. Interestingly, MIST1 also displays high structural homology with the CNL ZAR1 [72], predicted with 100% confidence over 700 amino acids (120-820) mostly outside MIST1 TIR domain, despite ZAR1 and MIST1 only sharing 18% sequence identity over the same region. It has been recently shown that a complex formed by ZAR1-RKS1-PBL2^UMP^ assembles into a high-order oligomeric complex in the form of a wheel-like pentamer (the resistosome), a structure that is required for immune signaling. The assembly is mediated by all the structural domains of ZAR (CC-NB-ARC-LRR), is further stabilized by ATP, and undergoes fold-switching during ZAR1 activation [72]. Considering the structural homology, it is tempting to speculate that MIST1 might also be assembling into high-order complexes *via* TIR-TIR oligomerization further stabilized by interaction among its NB-ARC-LRR domains, in a ZAR1-like manner. ZAR1 appears to be a recognition hub for at least three different bacterial effectors from different bacteria [73–75]. It might be the case that MIST1, on top of its function as a key regulatory hub for TNL expression at the RNA level, could also be acting as a recognition hub at the protein level.

*MIST1* is the first locus described in *Arabidopsis* that gives raise to phasiTNLs and, remarkably, triggers a larger accumulation of siRNA than any other NLR within the *Arabidopsis* genome (**Fig. 4**). This includes *RPS5*, *RSG1* and *AT5G43740*, the CNL gene targets of the RDR6-mediated regulation triggered by miR472 processing [25,34,52]. The finding that phasiRNAs generated from *MIST1* are loaded onto AGO1/ AGO2 complexes (**Fig. 8**), and that miR825-5p target site on *MIST1* can trigger *in trans* silencing of an AGAMOUS-based transitivity reporter (**Fig. 9**), supports the notion of these phasiRNAs being active and capable of establishing a second layer of regulation upon TNL gene expression. In further support of this notion, the predicted regulatory network of miR825-5p, including the five phasiRNA that accumulate from *MIST1* and are loaded onto AGO1 and/or AGO2 to highest levels, points to miR825-5p and its target *MIST1* as a key central hub for direct and indirect phasiRNA-mediated regulation of a very large number of TNLs. Indeed, when using default parameters for network prediction, the number of elements included in the miR825-5p/phasiTNLs network was unwieldy (**Table. S2**). This led us to apply more stringent parameters to reduce the size of the network to a manageable size (**Fig. 11**). We named <u>pha</u>siTNL-targeted <u>T</u>NLs (*PHATT* genes) those TNL genes within the network not directly regulated by miR825-5p but targeted by the phasiTNLs generated from *MIST1*. We found that phasiTNL4 displays a distinctly larger numbers of: (*i*) reads associated to AGO2 complexes (**Fig. 8**), and (*ii*) predicted target genes, seemingly acting as a secondary hub for TNL regulation. Interestingly, mapping of the phasiTNL4 target site onto the corresponding *PHATT4* genes showed this phasiTNL targets another highly conserved motif within the TIR domain, the TIR3 motif [42]. Thus, our results indicate that miR825-5p is part of the mechanism by which RDR6 acts as a negative regulator of plant immunity (**Fig. 2**, **3 and 7**).

**Figure 11.**
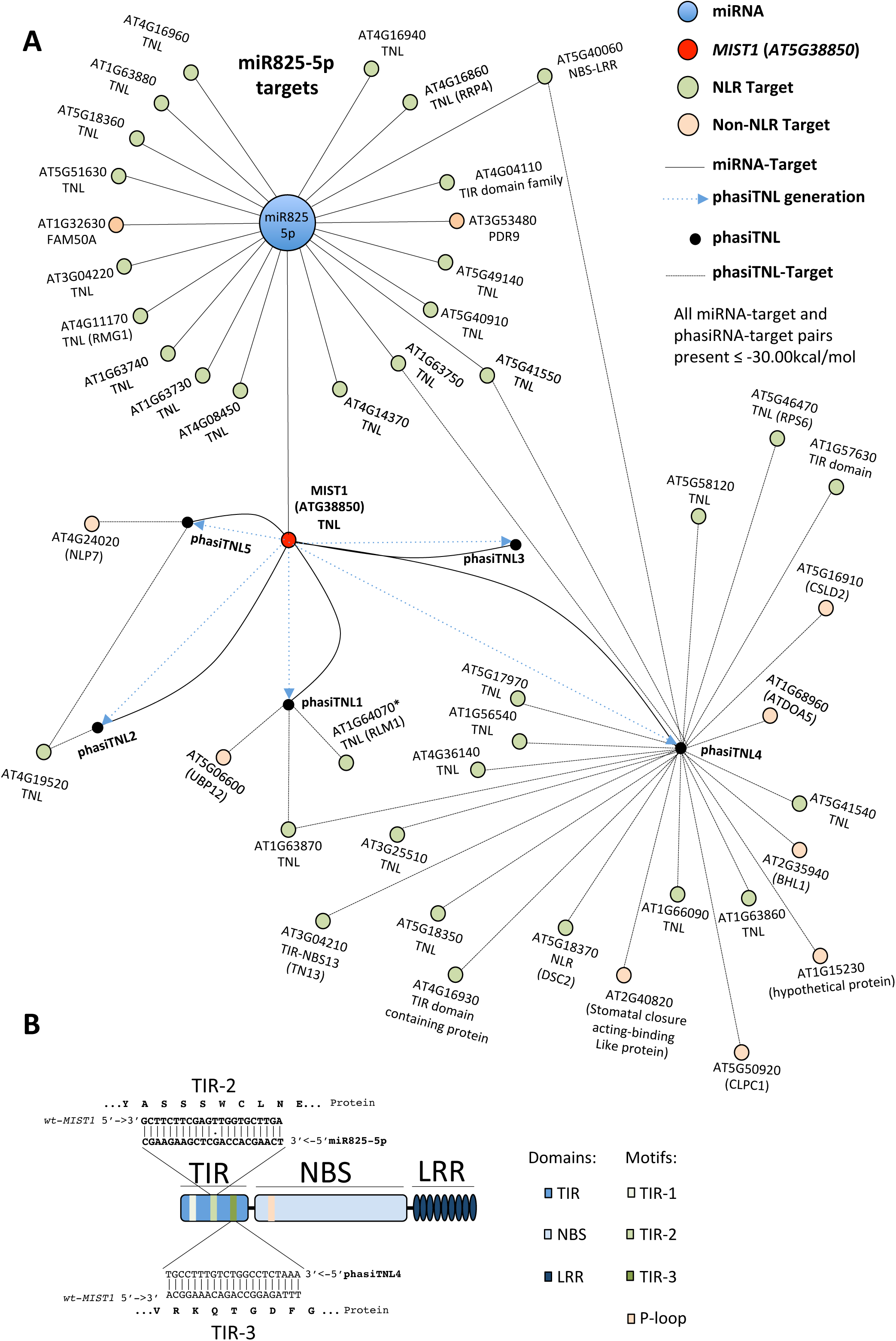
MiR825-5p predicted regulatory network. **A** MiR825-5p direct target genes are shown including MIST1 as a primary central hub for TNL gene regulation. Only the top five phasiTNLs in terms of accumulation and AGO1/AGO2 association are included, as well as their predicted targets. Hits of this phasiTNLs on primary miR825-5p are also indicated. PhasiTNL4 acts as a secondary hub in this regulatory network through targeting of the highly conserved TIR3 motif. **B** Domain organization of a TNL protein with domains and conserved motifs indicated. Alignments of miR825-5p and phasiTNLs with their respective target sequences are mapped to the corresponding domains/motifs within the TNL protein.

MiR472 and miR482/2118-mediated NLR silencing is lifted during the onset of PTI [33,34,36] when targeted NLR genes are likely to become transcriptionally active. In these circumstances, mature levels of these miRNAs decrease, lifting the silencing of NLR genes, while down regulation of RDR6 expression coupled with a reduction on siRNAs generated from these loci is expected to act by amplifying NLR gene activation. The data show that *MIR825* is transcriptionally down regulated during PTI and that this regulation is not restricted to response against flagellin, or for that matter against bacterial pathogens, since it also responds to the fungal PAMP chitin (**Fig. 10**), thus fitting into a model where miR825-5p silencing of TNLs would prevent the accidental onset of defenses in the absence of an incoming pathogen (**Fig. 12**). Interestingly, during preparation of this manuscript, a report established miR825-5p and 3p are involved in ISR triggered within the *B. cereus*/ *Botrytis cinerea* pathosystem [76]. In regards to the transcriptional down regulation of *MIR825*, a recent report showed that mutations that lead to over accumulation and nuclear localization of TNL protein SNC1 repress transcription of *MIR* genes, including *MIR825* [54]. These authors showed that *MIR* gene down regulation correlates with reduced accumulation of siRNA from several loci (including *MIST1*/*AT5G38850*), and increased expression of many NLR genes, and propose that SNC1 modulates immunity through the regulation of miRNA and phasiRNA biogenesis. Their results fit with those presented in here, opening the possibility of SNC1, likely together with co-repressor TPR1, being the mechanism responsible for *MIR825* transcriptional response to PAMPs. Taking all this into consideration, our model proposes that *in trans* silencing carried out by miR825-5p-triggered phasiTNLs generated from *MIST1* does not only play a role on amplifying the silencing signal on primary target TNL genes, but also by expanding the regulation to secondary targeted TNLs, different from those directly targeted by miR825-5p (**Fig. 11 and 12**). The results obtained by challenging with *P. syringae* transgenic plants displaying increased or reduced accumulation of miR825-5p (**Fig. 3**) demonstrate that miR825-5p mediated TNL silencing plays a relevant role in the regulation of plant immunity.

**Figure 12.**
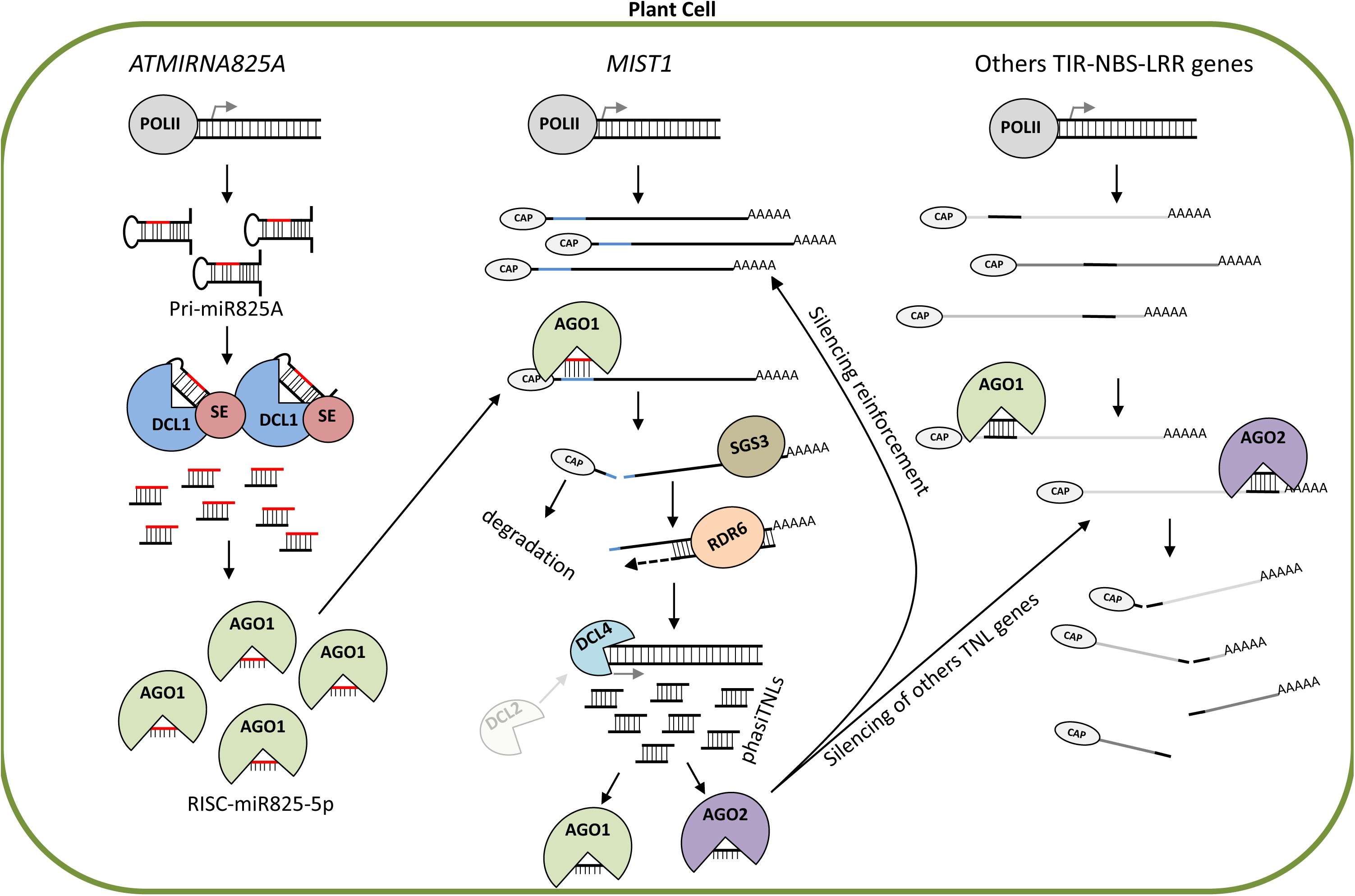
Model of the proposed regulatory network of miR825-5p. Under basal conditions, in the absence of a pathogen, miR825-5p silences expression of a number of TNL genes including *MIST1.* MiR825-5p silencing of MIST1 leads to the accumulation of numerous phasiTNLs through the action of RDR6 and DCL4/DCL2. These phasiTNLs act amplifying silencing of primary target TNL genes and silence in trans additional TNLs or PHATT (PHAsi-Targeted TNL) genes. Upon perception of PAMPs, perhaps involving SNC1, *MIR825* expression is down regulated, as has been reported for *RDR6*, leading to activation of TNL expression.

Targeting of conserved sequences encoding functionally relevant motifs provides a potential evolutionary link between protein function and miRNA-mediated regulation by coupling evolution of the one to evolution of the other. Indeed, such a link is also present in the miRNA-regulation of CNLs, and a few TNLs, in *M. sativa*. Most members of the miR472/miR482/miR2118 superfamily target the coding sequence for the highly conserved P-loop or Walker motif, essential for defense signaling [33], whereas miR1507 targets the Kinase-2 motif of the CC domain of a CNL-encoding gene [23]. No such evolutionary link has been established between function and regulation within the TIR domain prior to this report. MiR825-5p-mediated silencing of TNL is an example of coupled evolution of regulation of plant immunity, and the first that does so through targeting the coding sequence for a functional site within the TIR domain, the TIR2 motif, linked to the NAD+-cleavage enzymatic activity, thus specifically controlling TNL genes. Moreover, miR825-5p-dependent phasiTNL4 targets the sequence coding for TIR3, another highly conserved, motif within the TIR domain.

MiR825-5p is conserved in Brassicaceae (**Fig. 1;** [43, 46]). Using miR825-5p to search for potential target genes in the *B. oleareacea* genome in a recently created web browser (https://plantsmallrnagenes.science.psu.edu/; [77]), we found that several of its predicted targets, which display sequence similarity to *Arabidopsis* TNL genes, do accumulate significant amounts of 21 nt siRNAs, supporting the potential conservation of miR825-5p TNL regulatory network among Brassicaceae (**Fig. S3**). A recent report has identified an sRNA matching miR825-5p sequence as produced outside brassica species, in wheat (*Triticum aestivum L.*) [44]. Interestingly, wheat does not encode TNLs, since all its NLRs belong to the CNL family. TNLs have been proposed to arise in early plants and to have been later lost in monocots [43]. Since targets TNLs through TIR2, linked to cleavage of NAD+ cleaving activity [40, 67], miR825-5p might target genes encoding NAD+-cleaving enzymes in wheat. In this regard, a structure-based phylogeny inferred by [67] suggests that TIR domains might be part of a superfamily of structurally related proteins not usually associated to TIR domains (such as glycosyltransferases, nucleoside hydrolases or flavodoxins), but able to bind nucleotides in the analogous region of the protein.

Previous reports on miRNA-mediated regulation of CNLs have highlighted the role in lowering the cost of surveillance of miRNA-mediated regulation of CNLs, but these have also shown these miRNAs have a potential role in lowering the cost of defense [34, 35]. Our results support miR825-5p is involved in repressing TNL expression in basal, unchallenged, conditions and potentially be thus involved in lowering the cost of surveillance. Although no apparent fitness defect phenotype has been observed in plants with decreased miR825-5p accumulation, the fitness cost of these knock down lines have not been investigated in depth. In addition, our results also support miR825-5p role in keeping defenses under control during the onset of PTI. In this regard, it is interesting that miR825-5p regulation of TNLs, like the majority of miRNA-mediated regulatory networks involving NLRs described to date [34–36], has an impact on plant basal immunity, in the absence of R-mediated effector-recognition. NLR genes are normally associated with dominant gene resistance and this role was indeed the first described for a NLR gene [78], constituting the basis of race-specific resistance and involving the direct or indirect recognition of a pathogen effector protein. Although we cannot rule out an additional role for miR825-5p in regulating ETI, since miR825-5p targeted genes encode uncharacterized TNLs, it would have to be on the regulation of ETI mediated by a yet uncharacterized effector/R-gene pair in the *Arabidopsis*/ *P. syringae* pathosystem or by effectors from a different bacterial or fungal pathogen. On this note, AT5G18360, a putative primary target for miR825-5p, has been recently identified as a TNL involved in triggering ETI against Pto DC3000 effector HopB [79]. As proposed in tomato [35], miR825-5p may have a quantitative contribution to plant defenses, perhaps controlling low-level recognition of pathogen effectors in the absence of proper ETI. Further work will be necessary to characterize the role of the numerous TNL genes belonging to the miR825-5p regulon.

## Methods

### Plant material and growth

*Arabidopsis thaliana* plants were grown in soil, or MS plates without sucrose, at 21°C with a photoperiod of 8h light/16h darkness with a light intensity of 200 µmol/m^2^/s. For MS growth, seeds were surface sterilized with a mixture of ethanol and bleach for 15 min and washed three times with ethanol. In all cases seeds were stratified for 2 days at 4°C. When selecting transformants, MS plates supplemented with either kanamycin (50 μg/ml) or hygromycin (40 μg/ml) were used. *Nicotiana benthamiana* plants were grown in soil in temperature-controlled chambers, at 22°C with a controlled photoperiod of 16h light/8h dark with a light intensity of 200 µmol/m^2^/s.

### DNA procedures

All DNA fragments generated by PCR for cloning were amplified using the Q5 High-Fidelity DNA polymerase (NEB, USA), as indicated by manufacturer. Depending on the construct, these fragments were alternatively cloned into pGEM-T (Promega, USA) or into pENTR (Themo Fisher Scientific, USA) digested with *Not*I-*As*CI. Derivatives of pGEM-T thus obtained were further digested to obtain the PCR-amplified cloned fragments for classic ligase-based cloning into the corresponding destination vector. For PCR-amplified fragments initially cloned into pENTR, LR reactions were carried out to generate the final vectors. For a detailed description on the cloning process for each specific construct see supplemental material. Routine PCRs for cloning confirmation or plant genotyping were performed using Gotaq Flexi DNA Polymerase (Promega, USA) following instructions from the manufacturer. Sequences for primers used in this work can be found in **Table S3**.

### RNA procedures

Total RNA from seedlings or adult leaves was extracted using TRISURE (Bioline, UK) according to instructions from the manufacturer, using 100 mg of the corresponding plant tissue, previously frozen and grounded in liquid nitrogen. RT reactions were performed using iScript cDNA Synthesis Kit (Bio-Rad, USA) and 1 µg of total RNA, previously treated with DNaseI (Takara, Japan).

For semi-quantitative PCRs quantification of pri-miR825, we used 1 µl of the cDNA obtained as described, and the number of cycles was restricted to non-saturating conditions (typically 20-28 cycles depending of the gene).

For RT-*q*PCRs, we used 2 µl of a 1/5 dilution of the cDNA obtained as above in a reaction containing 5 µl of SsoFast EvaGreen (Bio-Rad, USA), 0.5 µl of Forward primer (10 µM) 0.5 µl Reverse primer (10 µM) and 2 µl of H2O. These RT-*q*PCRs were performed using a CFX96 or CFX384 machine (Bio-Rad) with a first denaturing step at 95°C 1 min, and 45 cycles of 95°C 10 s and 60°C 15 s. In all cases ACT2 was used as internal control. Relative expression was calculated using the 2^−ΔΔCt^ method [80].

For quantification of mature miRNAs, we used the stem-loop RT-*q*PCR method [81]. Pulsed-RT (1 step at 16°C of 30 min, followed by 60 cycles at 30°C for 30 s, 42°C for 30 s and 50°C for 1 min, and 1 cycle of 85°C for 5 min) was performed using Revert Aid First Strand cDNA Synthesis Kit (Thermo Fisher Scientific, USA) with specific RT stem-loop primers and oligo (dT). Stem loop RT-*q*PCRs were performed in a CFX96 or CFX384 (Bio-Rad) machine using the protocol previously described by[81].

For small RNA northern blot analysis, we extracted total RNA using TRISURE (Bioline, UK), and use it for Northern blots analyses carried out as previously described by [82]. In brief, 50 μg of total RNA were suspended into 2x RNA loading buffer (95% formamide, 18 mM EDTA pH 8.0, 0.025% sodium dodecyl sulfate (SDS), 0.01% bromophenol blue, 0.01% xylene cyanol) and denaturized at 90°C for 5 min. Then, samples were separated by electrophoresis in a 7M urea, 0.5X TBE, 17% polyacrylamide gel. After that, RNA was electro-blotted onto an Amersham Hybond-N+ nylon membrane (GE Healthcare Life Sciences, USA) at 80 V for 1 hour in cold 0.5x TBE. RNA was fixed onto the membrane by exposing it to, 0.120 J of UV after which the membrane incubated at 80°C for 1 hour. Afterwards, the membrane was pre-hybridize in Church buffer (1% BSA, 1 mM EDTA, 0.5 M phosphate buffer, 7% SDS) for 1 h at 40°C. For hybridization, the probe was added to Church buffer and incubated overnight at 40°C. The next day, membranes were washed 3 times with a 2x SSC, 0.1% SDS solution at 40°C (10 min per wash) and detection was carried out as previously described by [82].

For probe labelling, a DNA oligonucleotide reverse complement to U6 was 3’-end-labelled with Digoxigening-11-ddUTP (Sigma, USA) using a Terminal Deoxynucleotidyl Transferase (TdT; ThermoFisher SCIENTIFIC, USA) in a reaction containing: 20 Units TdT, 10 μl 5x TdT reaction buffer, 5 μl DNA primer (1 μM), 2.5 μl Digoxigenin-11-ddUTP (10 μM) and bidistilled H_2_O to 50 μl. The reaction was incubated for 40 min at 37°C and directly added to the hybridization solution without further purification.

For secondary siRNA detection, a fragment of *MIST1* was PCR amplified using Q5 DNA polymerase (NEB USA) with MIST1_PhasiRNA_ProbeF and MIST1_PhasiRNA_ProbeR. PCR product was gel-purified and 400 ng of the purified product were used in a random priming reaction containing: 4 Units Klenow fragment (TAKARA, Japan), 5 μl 10x Klenow reaction buffer, 12 μl Random hexamers (100 μM), 5 μl of 10x DIG DNA labelling Mix (Sigma, USA) and H2O to 50 μl.

### Protein extraction and Western blot

Approximately 100 µg of leaf tissue were harvested, frozen into liquid nitrogen and grounded into 100 µl of extraction Laemmli buffer (62.5 mM Tris-HCl pH 7.4, 100 mM Dithiothreitol (DTT), 2% sodium-dodecyl sulfate (SDS), 0,001% Bromophenol blue (BPB), and 10% glycerol). The resulting homogenate was centrifuged at 20000 *g* for 10 min at 4°C. Soluble supernatant was centrifuged again in a fresh tube to ensure absence of insoluble debris. Protein concentration was determined by the Bio-Rad protein assay (Bio-Rad, USA). Ten micrograms of each protein sample, unless otherwise stated, were resolved on 10-12% acrylamide SDS-PAGE gels (Mini protean, Bio-Rad, USA) and transferred onto nitrocellulose membranes (Immobilon-P, Millipore, USA), using the Semi-Dry Transfer System (Bio-Rad, USA) during 1h at 25V. Western blots for immunodetection of GFP (Santa Cruz Biotechnology, USA), Tubulin (Abiocode, USA), MPKs (Cell Signaling Biotechnology, USA), or anti-PR [83] were carried out using standard methods, with a 1:600 dilution of primary anti-GFP, 1:1000 for anti-Tubulin, 1:5000 for anti-MPKs and 1:5000 for anti-PR1. For secondary antibodies, 1:10000 dilution of a secondary Anti-Rabbit antibody (SIGMA, USA), and 1:80000 dilution for Anti-Mouse antibody (SIGMA, USA) were used. Membranes were developed using the Bio-Rad Clarity Western ECL Substrate (Bio-Rad, USA) following instructions from the manufacturer.

### Bacterial assays

Bacterial *in planta* assays were carried out using *P. syringae pv.* tomato DC3000 [84] or a derivative carrying a plasmid constitutively expressing avirulence effector AvrPt2 [85]. Colonies from Lysogenic Broth (LB) medium [86] plates incubated for 2 days at 28°C. Bacterial plant inoculations for RT-*q*PCR or semi-quantitative PCR analysis of inoculated leaf tissue were carried out using a 10 mM MgCl_2_ bacterial suspension at 5×10^7^ colony forming unit per ml (cfu/ml) to pressure-infiltrate *Arabidopsis* adult leaves using a needleless syringe. Inoculations for bacterial proliferation assays were carried out using bacterial colonies obtained as above but infiltrating leaves with bacterial suspensions at 5×10^4^ cfu/ml. In these assays, 4 days post inoculation (dpi) three inoculated leaves per plant were collected and one leaf disk of 10 mm diameter obtained from each that were grinded together into 10 mM MgCl_2_ to generate a biological replicate. Serial dilutions were then plated onto LB plates (supplemented with cycloheximide at 2 µg/ml to prevent fungal contamination), which were incubated for 2 days at 28°C to calculate cfu/cm^2^.

For transient expression assays in *N. benthamiana*, 4-5 weeks old plants were infiltrated with an *Agrobacterium tumefaciens* (GV3101 or C58C1 strains) [87] carrying the corresponding binary plasmids (**Table S4**). Inoculations were carried out using saturated LB 28°C cultures (and the corresponding antibiotic at the following concentration: 50 μg/ml kanamycin, 50 μg/ml rifampicin, 20 μg/ml gentamycin, or 5 μg/ml tetracycline), diluted into infiltration medium (10 mM MES (SIGMA, USA), 10 mM MgCl_2_, 150 µM 3′,5′-Dimethoxy-4′-hydroxyacetophenone (acetosyringone; SIGMA, USA) at a 23.7×10^8^ cfu/ml (OD_600_=0.75) or infiltration medium (mock controls). Samples were taken 2 to 3 days post-inoculation.

### Generating *Arabidopsis* transgenic lines

To generate transgenic lines (**Table S5**), 5 mL of LB inoculated with *A. tumefaciens* was incubated overnight at 28°C with agitation, and 500 µL of it used to inoculate 100 mL of LB supplemented with the corresponding antibiotic that was also incubated at 28°C with agitation for 24 h. The resulting *A. tumefaciens* culture was centrifuged at 5500 *g* for 10 min. After discarding the supernatant, pellets were suspended into 50 mL of H_2_O containing sucrose (50 g/L) and Silwet (50 µl/L; UNIROYAL CHEMICAL, UK). These suspensions were used for floral dipping to generate the *A. thaliana* transgenic lines [88]. Putative transformants were selected into plates of MS supplemented with Km (50 μg/ml) or hygromycin (40 μg/ml), and the presence of the transgen confirmed by PCR as described above.

### Eliciting basal plant defense responses

To elicit basal plant responses, either a 100 nM solution of flg22 immunogenic flagellin peptide (GenScript), or a 0.2% solution of chitin (method2 from [89]), were needless syringe-infiltrated into adult plant leaves. Each assay included plants infiltrated with water as mock treated control, to discard differences observed were caused by pressure infiltration-associated mechanical damage. All infiltrated tissues were harvested, immediately frozen in liquid nitrogen at the indicated time points after treatment, and stored at −80°C until their use in the subsequent assays.

### MAPK activation assays

For MAPK activation assays, *A. thaliana* seedlings (4 per sample) were grown in MS plates at 21°C with a photoperiod of 8h light/16h darkness and a light intensity of 200 µmol/m^2^/s. Twelve-days-old seedlings were transferred into liquid MS and maintained there for 24 h. Seedlings were then transferred into 12-well plates containing a 100 nM solution of flg22, and samples were collected and frozen into liquid nitrogen at the indicated times. Frozen samples were grounded and proteins extracted in a buffer containing 100 mM Tris-HCl, 150 mM NaCl and 1x Halt Phosphatase Inhibitor Cocktail (Thermo Fisher Scientific, USA). Extracted proteins were mixed with a 3x Laemmli buffer (2% SDS, 10% glycerol, 100 mM DTT, 0.001% BPB and 0.0625 M Tris-HCL) loaded and separated into a 12% SDS-PAGE, and then were transferred onto nitrocellulose membrane as describe above. Finally, membranes were incubated with the antibody against Phospho-p44/42 MAPK (Erk1/2) (Thr202/Tyr204) (D13.14.4E) XP® (Cell Signaling Technology, USA). As a loading control membranes were incubated with anti-tubulin antibody (Abiocode, USA).

### ROS quantification

*Arabidopsis* plants of the indicated genotypes were grown 2-3 weeks on soil during 2-3 weeks as described above. Two leaf disks were taken per plant with a cork borer (diameter=3.8 mm), transferred into a 96-well plate containing 100 µl of water, and incubated for 24 hours at room temperature. The next day, water was removed and replaced by 100 µl of the assay solution: 17 µg/mL Luminal (Sigma-A8511, USA), 10 µg/ml Horseradish peroxidase (HRP) (Sigma-P6782, USA), 100 nM flg22 (see above) and water. Light emission was measured immediately in a GloMax 96 Microplate Luminometer (Promega, USA). At least 16 leaf disks were taken by treatment (n ≥ 16).

### Bioinformatic analysis

Raw files were obtained from Sequence Read Archive (NCBI). These files were converted to fastq files using the SRA toolkit (fastq-dump; http://ncbi.github.io/sra-tools/), and quality filtered with removal of adapter content using Trimmomatic [90]. Afterwards, quality was confirmed using FASTQC (Simon Andrews, bioinformatics.babraham.ac.uk/projects/fastqc/), and the reads mapped against the *A. thaliana* genome (TAIR 10) using Bowtie [91] with no mismatch allowed (–v 0 mode) (**Fig. 4B and 7**), except for AGO1 and AGO2 libraries (-v 1 mode) (**Fig. 1E and Fig. 8B-D**). SAM files were converted to BAM, sorted and indexed using samtools [92]. Mapping of the reads was visualized using IGV browser [93]. Numbers of reads mapped per feature of the *A. thaliana* genome were estimated using HTSeq (htseq-count; [94]) and normalized as number of reads mapping to the feature per ten millions of total reads mapped to the entire genome (RPTM method). The AGO1 and AGO2 IP libraries used in this study (**Fig. 1E and Fig. 8**) correspond to: GSM2787769, GSM2787770, GSM3909547 and GSM3909548. For **Fig. 4B and 7** the raw reads were retrieved from SRP097592 bioproject.

MicroRNA and siRNA target prediction (**Fig. 1A**, **Fig. 4A**, **Fig. 11, Table S1 and Table S2**) was performed using WMD3 (http://wmd3.weigelworld.org/cgi-bin/webapp.cgi) and/or psRNATarget ([41]) with default parameters. To analyze miRNA825 conservation across different species (**Fig. 1C**), sequences were retrieved from miRBase [95] or from NCBI (Blastn against the desired genomes with ath-miR825 as a template), aligned using Clustal Omega [96], and logos were generated using Weblogo [97]. NLRs Protein sequences used in Figure 1B were retrieved from TAIR. The logo was regenerated as described above.

To determine miR825 5p/3p ratios (**Fig. 1D**), raw files were obtained from Sequence Read Archive (NCBI) under the accession numbers: (SRR2079799, SRR2079800, SRR2079801, SRR2079802, SRR2079803, SRR2079804, SRR2079805, SRR2079806, SRR2079807, SRR2079808, SRR2079809, SRR2079810, SRR2079811, SRR2079812) and converted to fastq files as explain above. Then, the number of 5p and 3p reads were investigated using the “grep” command. Only reads starting with mature miRNA sequences and containing the adapter sequence immediately after miRNA reads were used for further analysis. Ratios were calculated for each library, and represented as miR825-5p/miR825-3p.

For Fig. **6A**, secondary RNA prediction was carried out using Mfold (http://unafold.rna.albany.edu/?q=mfold) and the precursor was visualized with Varna (http://varna.lri.fr/index.php?lang=en&page=home&css=varna). Secondary and tertiary structures for the miR8255-p/miR825-3p duplex was predicted using Mfold and MC-fold/MC-Sym as previously described [53].

## Acknowledgments

We are very grateful to Adela Zumaquero whose work inspired the work reported here. We also wish to thank Pablo García Vallejo for technical assistance. This work was supported by project grants from Ministerio de Economia y Competitividad (MINECO, Spain; BIO2015-64391-R) and Ministerio de Ciencia, Innovación y Universidades (MCIU, Spain, RTI2018-095069-B-100) awarded to C.R. Beuzón and J. Ruiz-Albert, and Junta de Andalucía Proyecto Operativo FEDER Andalucía (Spain; UMA18-FEDERJA-070) to J. Ruiz-Albert and E.R. Bejarano. The work was co-funded by Fondos Europeos de Desarrollo Regional (FEDER). D. López-Márquez was supported by an FPU Grant (Predoctoral Fellowship from the Spanish Ministerio de Educación, Cultura y Deporte; FPU14/04233), Plan Propio Universidad de Málaga (UMA) and project grant RTI2018-095069-B-100 awarded to C.R. Beuzón and J.Ruiz-Albert. ADE was funded by a FPU Grant (Predoctoral Fellowship from the Spanish Ministerio de Ciencia, Innovación y Universidades; FPU17/03520). The work carried out in Ignacio Rubio-Somozás laboratory was funded by FEDER/Agencia Estatal de Investigación (AEI)/ Ministerio de Ciencia, Innovación y Universidades (MCIU, Spain, RTI2018-097262-B-I00)/Ministerio de Economía y Competitividad (MINECO, Spain, RYC-2015-19154 and BFU2014-58361-JIN), with the financial support from the Ministerio de Economía y Competitividad (Severo Ochoa Programme for Centres of Excellence in R&D 2016-2019, SEV-2015-0533) and the CERCA Programme/Generalitat de Catalunya.

## Author Contributions

DLM, ERN, IR-S, JRA, ERB and CRB contributed to the conception of the work and the experimental design. Data acquisition and primary analysis has been the result of the work of DLM, ADP, NLP, ERN and JRA, while ALL authors participated into data interpretation. The paper has been drafted by the combined efforts of DLM, JRA, ERB and CRB, with the final version settled by ALL authors after critical revision. ALL authors approved the final submitted version, and agree to be accountable for the accuracy and integrity of their respective contributions to the work presented in this paper.

## Figure legends

**Figure S1. SiRNA production from different NLR genes. Images show** screenshots from MPSS showing sRNAs that originate from NLR genes in *Arabidopsis* selected among those displayed in Fig. 4B for accumulating the highest levels of sRNAs.

**Figure S2: *A*tmiR825A promoter is active in adult leaves. A.** Confocal microscopy images showing GFP accumulation in a transgenic line harbouring the reporter gene under the control of *At*miR825 promoter (pMIR825A::GFP-HIS). **B**. Western blot analysis using anti-HIS antibody show accumulation of the GFP-HIS reporter fusion protein in several transgenic lines harbouring the pMIR825A::GFP-HIS construct. The membrane was stained with Coomassie and used as loading control. Samples for **A** and **B** were taken from *Arabidopsis* adult leaves.

**Figure S3: Predicted targets of miR825-5p in *Brassica oleracea*. Upper panels of A and B:** Screenshots showing the complementarity between miR825-5p and the potential targets predicted using psRNATarget. **Lower panels of A and B:** Screenshots showing the accumulation of 21/22-nt sRNAs from these potential targets. The most similar gene in *Arabidopsis* is indicated.

**Table S1. Extended list of targets for miR825-5p and 3p using two different prediction software.**

**Table S2. Extended list of primary and secondary targets of the miR825-5p/MIST1/phasiTNLs predicted network (including a larger number of phasiTNLS and standard parameters)**

**Table S3. Primers used in this work.**

**Table S4. Plasmids used in this work.**

**Table S5. Transgenic lines generated in this work.**

